# Convergent and lineage-specific genomic changes shape adaptations in sugar-consuming birds

**DOI:** 10.1101/2024.08.30.610474

**Authors:** Ekaterina Osipova, Meng-Ching Ko, Konstantin M. Petricek, Simon Yung Wa Sin, Thomas Brown, Sylke Winkler, Martin Pippel, Julia Jarrells, Susanne Weiche, Mai-Britt Mosbech, Fanny Taborsak-Lines, Chuan Wang, Orlando Contreras-Lopez, Remi-Andre Olsen, Philip Ewels, Daniel Mendez-Aranda, Andrea Gaede, Keren Sadanandan, Gabriel Weijie Low, Amanda Monte, Ninon Ballerstaedt, Nicolas M. Adreani, Lucia Mentesana, Auguste von Bayern, Alejandro Rico-Guevara, Scott V. Edwards, Carolina Frankl-Vilches, Heiner Kuhl, Antje Bakker, Manfred Gahr, Douglas L. Altshuler, William A. Buttemer, Michael Schupp, Maude W. Baldwin, Michael Hiller, Timothy B. Sackton

## Abstract

High-sugar diets cause human metabolic diseases, yet several bird lineages convergently adapted to feeding on sugar-rich nectar or fruits. We investigated the underlying molecular mechanisms in hummingbirds, parrots, honeyeaters, and sunbirds by generating nine new genomes and 90 tissue-specific transcriptomes. Comparative screens revealed an excess of repeated selection in both protein-coding and regulatory sequences in sugar-feeding birds, suggesting reuse of genetic elements. Sequence or expression changes in sugar-feeders affect genes involved in blood pressure regulation, lipid, amino acid and carbohydrate metabolism, with experiments showing functional changes in honeyeater hexokinase 3. *MLXIPL*, a key regulator of sugar and lipid homeostasis, showed convergent sequence and regulatory changes across all sugar-feeding clades; experiments revealed enhanced sugar-induced transcriptional activity of hummingbird *MLXIPL*, highlighting its adaptive role in high-sugar diets.

**Summary Figure: Our comparative screens across four independent sugar-feeding bird groups identified both repeated and lineage-specific targets of selection:** 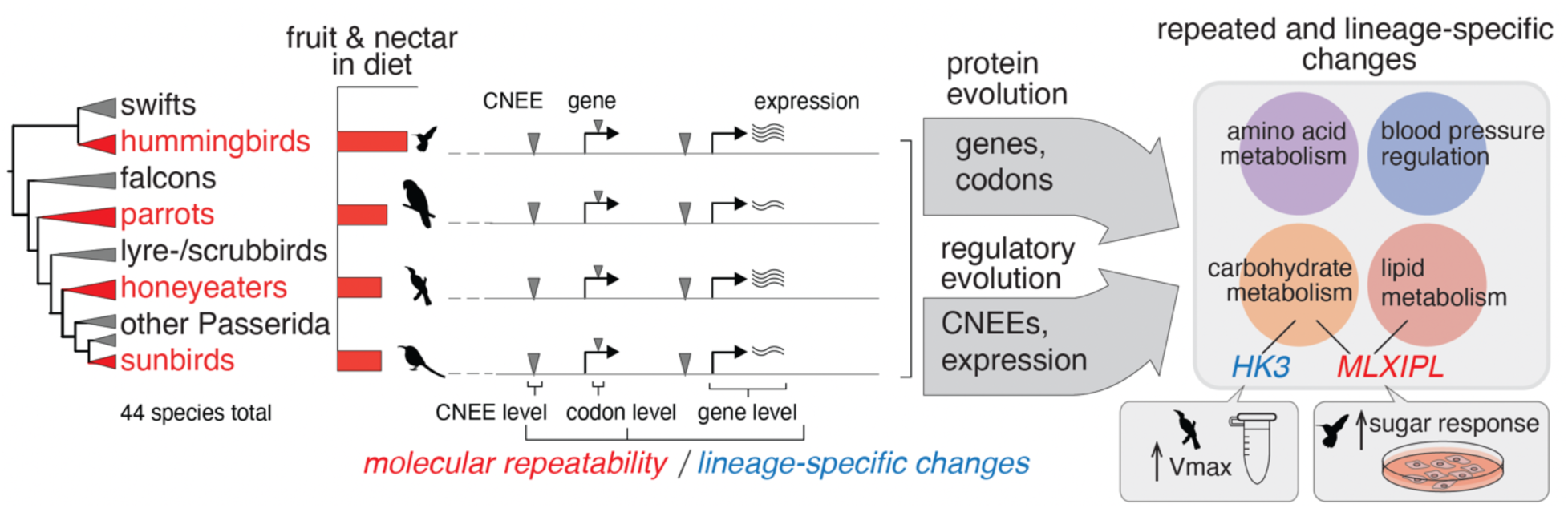

---

Animals vary tremendously in the foods they consume, and this imposes diverse requirements on their morphology, physiology, and behavior. Among birds, several groups including hummingbirds, sunbirds, and many honeyeaters and parrots independently adapted to feeding primarily on sugar-rich nectars and fruit (*1*, *2*). Nectar provides mostly sugars (*3*), primarily sucrose, glucose, and fructose (*4*), but contains little protein and fat, a challenge that birds overcome by supplementing nectar consumption with insects or pollen (*5*). In humans, excessive sugar consumption is a major risk factor for type 2 diabetes mellitus (T2D), a disease characterized by the failure of organs to regulate glucose uptake and production in response to insulin (insulin resistance) (*6*). In contrast, nectar-feeding birds have evolved physiological mechanisms to tolerate high-sugar diets without developing metabolic disorders. Understanding the morphological and metabolic adaptations that enable nectarivorous birds to thrive on their sugar-rich diet may provide insights that could be ultimately harnessed for human metabolic disease therapy.

Adaptation to high-sugar diets in birds occurs in the context of several important physiological differences compared to mammals. Birds naturally sustain blood glucose levels that are 1.5-2 times higher than mammals of similar body size (*7*, *8*). This hyperglycemia is partly attributed to insulin insensitivity linked to the lack of the glucose transporter GLUT4, which drives insulin-mediated glucose uptake in mammals (*7–9*); although see (*10*). Instead, birds rely on constitutively expressed glucose transporters, such as GLUT1 for systemic sugar uptake (although the use of these transporters may also vary by species) (*11*). Furthermore, in contrast to mammals, glucagon plays a dominant role in avian glycemic control, with high expression and constitutive activity of the glucagon receptor driving high blood glucose levels, potentially impacting lipid metabolism and contributing to the high metabolic rates observed in birds (*12*).

In birds, nectarivory has evolved multiple times (*13*, *14*), and nectar-feeding lineages share several common adaptations to their sugar-rich diet. Morphologically, they share elongated bills and specialized tongues that facilitate nectar extraction (*15–17*). At the physiological level, nectar specialists show elevated sucrase activities for efficient sucrose hydrolysis, with hummingbirds, honeyeaters, and sunbirds exhibiting significantly higher activities than non-nectar feeders, while lorikeets show more modest increases (*18*). These birds also possess mechanisms to manage the substantial water loads from nectar consumption, through adaptations in kidney function (hummingbirds) and, in some cases, modified intestinal water handling (honeyeaters and sunbirds) (*19–22*). Despite differences in strategies across lineages, shared physiological challenges likely drove selection on functional pathways that allow them to cope with high sugar loads and osmotic stress.

Previous studies, largely focused on hummingbirds, have identified some genomic correlates of physiological and metabolic adaptations to high-sugar diets. For example, pseudogenization of the glyconeogenic muscle enzyme FBP2 in hummingbirds enhanced glycolysis and mitochondrial respiration in muscle cells and likely facilitated the evolution of their energy-demanding hovering flight (*23*). Transcriptomic and genomic analyses of hummingbirds have suggested fast evolution of lipogenic pathways (*24*) and revealed changes in sugar transporters (*25*). Studies of taste receptors have revealed that hummingbirds and songbirds have convergently repurposed the amino-acid taste receptor (T1R1-T1R3) to sense sugars in nectar (*26*, *27*). Despite these advances, two main research gaps persist. First, many relevant genomic adaptations remain to be discovered. Second, the lack of systematic studies examining multiple nectar- or fruit-feeding lineages limits our understanding whether convergent sugar diet adaptations involve changes affecting the same gene across independent clades, lineage-specific changes affecting distinct (but potentially functionally-similar) genes, or both. Repeated evolution - defined here as shared changes or evolutionary patterns affecting the sequence or regulation of the same gene in at least two independent lineages - can indicate constrained possibilities for adapting to a high-sugar diet. The multiple independent transitions of bird lineages to sugar-rich diets thus provides a powerful system to investigate the role of repeated evolution at the molecular and sequence level in adaptation, a central question in evolutionary biology.

To address both gaps, we combined genome sequencing with comprehensive comparative genomic and transcriptomic analyses and experimental validations to uncover both repeated and lineage-specific genomic adaptations to a sugar-rich diet in four avian clades containing nectar-specialists: hummingbirds, sunbirds, honeyeaters, and parrots. We include species with high levels of fruit or nectar consumption in these four focal ‘sugar-consuming’ lineages. Whereas in hummingbirds, elevated sugar consumption evolved once and was retained throughout the clade, in songbirds and parrots the evolutionary history of sugar sensing is more complex, as sugar may have been an early part of the diet, with specialist nectarivore lineages evolving multiple times (*13*, *27*). Nevertheless, high levels of sugar intake in honeyeaters, sunbirds, and many parrots likely led to elaborated metabolic and physiological adaptations, detectable as genomic signatures of sequence change. We present an integrated analysis of several types of evolutionary signatures that can indicate a shift in function, considering protein-coding genes, gene expression, as well as non-coding genomic regions, to address the prevalence of repeated or lineage-specific molecular changes at multiple levels.

## Genome assembly, annotation, and comparative analyses

To investigate the genomic basis of adaptations to sugar-rich diets across hummingbirds, parrots, honeyeaters, and sunbirds, we used long and short read sequencing to assemble new reference genomes for five nectar- or fruit-taking birds and four outgroup species (Fig. 1A-C, table S1). These genomes are highly complete: we recovered 95.5% to 97.3% of 8,338 universal single-copy avian orthologs (BUSCO odb10 gene set) (*28*) (fig. S1C, table S1). We also assessed gene completeness with TOGA, a method that integrates comparative gene annotation, inferring and classifying orthologous genes (*29*), which recovered ≥10,092 intact coding genes of 15,133 ancestral bird genes across our nine new assemblies (Fig. 1D). Together with 35 previously generated assemblies (*30*, *31*), these genomes covered all four sugar-consuming clades, each with several representative species, and provided the basis for our analysis.

**Fig. 1.**
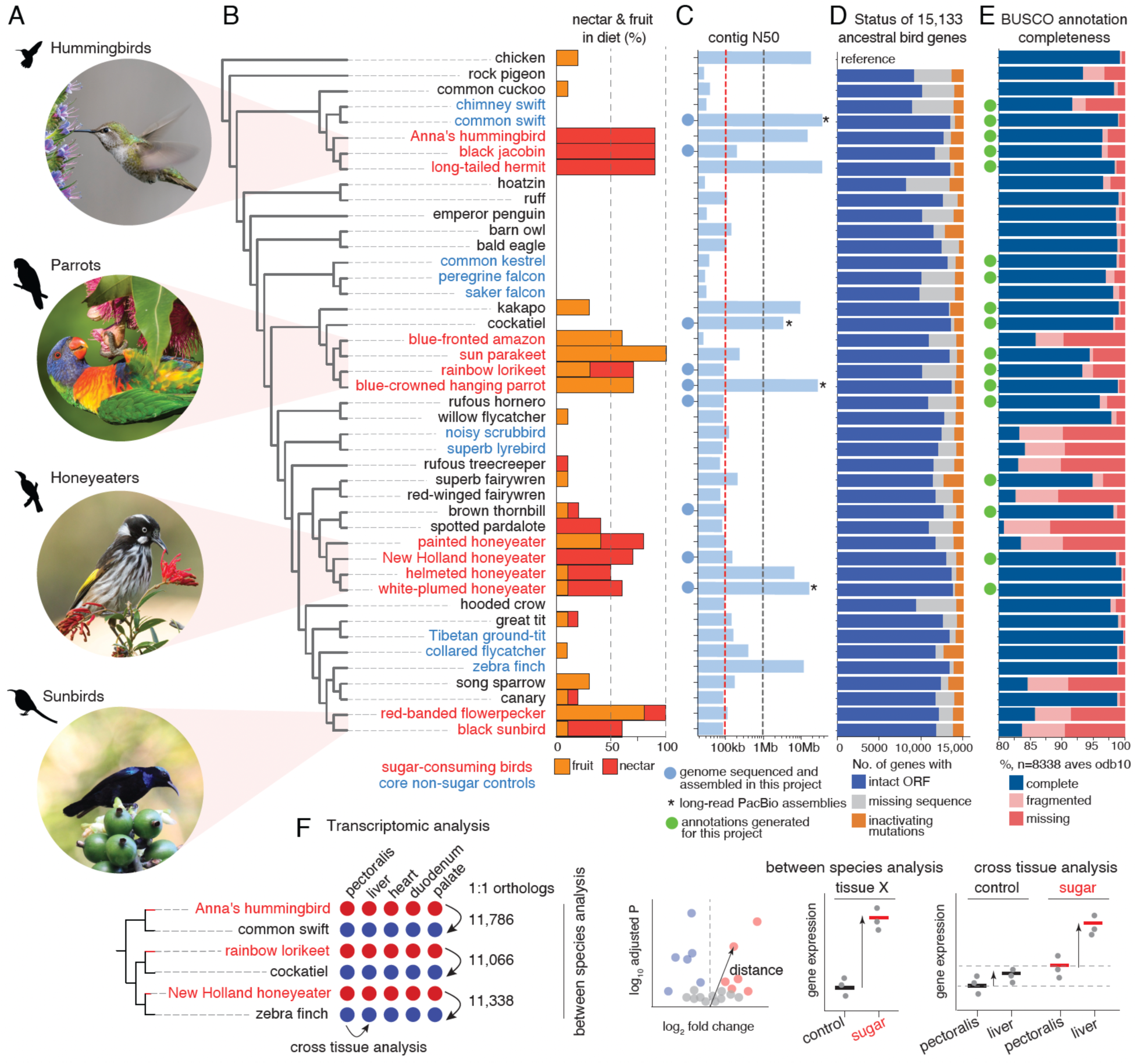
Genome assemblies, annotations, and transcriptomes. (A) Species representing the four sugar-consuming groups. Bird images from the Macaulay Library at the Cornell Lab of Ornithology: ML426078561, ML431374521, ML285318411, ML205719751. (B) Cladogram and diet of the birds included in the study. Based on the percentage of dietary nectar and fruit intake, we delineated four groups of core sugar-consuming birds (red font) and core non-sugar controls (blue). Core non-sugar control sets were defined as groups phylogenetically close to sugar-consuming clades and comprising a similar number of species, but not taking any nectar in the diet. These were used as direct controls in the downstream analyses. (C) Comparison of genome assembly contiguity. The contig N50 value indicates that 50% of the assembly consists of contigs of at least that size. The nine newly-sequenced birds are indicated with a blue circle. (D) Status of 15,133 ancestral bird genes, classified by TOGA as those with an intact reading frame (blue), with gene inactivating mutations (premature stop codons, frameshifts, splice site disruptions, and deletions of exons or entire genes; orange), or missing or incomplete coding sequences due to assembly gaps or fragmentation (gray). (E) Status of 8,338 near-universally conserved Aves genes (BUSCO, odb10) in annotations generated in this study (green circle) or NCBI RefSeq annotations. (F) Tissue-specific transcriptomes generated for pairs of sugar-consuming and non-sugar control species. Black arrow specifies the direction of DESeq2 analysis (for details, see Methods and table S11).

Comprehensive protein-coding gene annotations are key to studying molecular changes underlying phenotypic differences. To produce highly complete protein-coding gene annotations, we adapted a gene annotation pipeline (*32*) that combined TOGA’s ortholog predictions, transcriptomic data, protein alignments, and ab initio gene predictions (fig. S1D). Applying this pipeline to our nine new and eight previous assemblies that lacked an annotation generated by NCBI (*33*) produced annotations with a high BUSCO completeness of 93.2% - 99.6% for all 17 genomes, comparable or better than existing NCBI annotations (Fig. 1E).

Adaptations to sugar consumption may be the result of the repeated and lineage-specific evolution of protein-coding gene sequences, regulatory sequences, or gene expression (*23*, *25*). To investigate protein-coding evolution, we screened 11,688 one-to-one orthologs identified with TOGA (*34*) across 44 bird genomes for signatures of episodic positive selection in each branch in the phylogeny using the branch-site method implemented in aBSREL (*35*). We also identified convergent amino acid substitutions in sugar-consuming clades with CSUBST (*36*). To investigate regulatory evolution, we screened 363,476 avian conserved non-exonic elements (CNEEs) (*37–40*) – short, conserved sequences in introns or intergenic regions that represent putative enhancers – for rate shifts in one or more sugar-consuming lineage with the Bayesian method PhyloAcc (*41*). All these methods are designed to account for phylogenetic relationships by analyzing individual branches or branch pairs within the phylogenetic tree. Furthermore, we generated 90 total transcriptomic (RNA-seq) datasets for five tissues relevant to metabolism and digestion for three pairs of sugar-consuming species with a non-sugar outgroup, each in three biological replicates (Fig. 1F, table S9). We investigated gene expression changes with (i) phylogenetically independent pairwise differential expression tests between each sugar-consuming species and the respective non-sugar outgroup, considering comprehensive sets of >11,000 one-to-one orthologs and ranking expression changes by a metric that jointly considers significance and effect size (Fig. 1F); and (ii) a multifactor analysis conducted independently on the same pairs to detect smaller sugar-diet-associated tissue-specific expression shifts between the key metabolic tissues - pectoralis and liver - that are likely explained by selection rather than drift (see Fig. 1F and Methods). Together, these methodological approaches provide a comprehensive analysis of genomic and transcriptomic changes in sugar-consuming species.

## Sugar-feeding lineages have an excess of repeated selection in both protein-coding and regulatory sequences

We first addressed whether convergent sugar-consuming bird lineages exhibit more repeated evolution in genes and regulatory elements than matched control lineages. Our positive selection screen revealed 3,093 protein-coding genes selected on at least one branch in the 44-bird phylogeny, of which 592 (19%) are under selection specifically in sugar-consuming branches. These 592 genes include 254, 178, 109, and 126 selected on at least one branch in the hummingbird, honeyeater, parrot, and sunbird clades, respectively (Fig. 2A, table S2). Of these genes, 66 (11.1%) are under selection in multiple sugar-consuming lineages, with 58, seven and one genes being selected in two, three and all four lineages (Fig. 2A). To test whether repeated selection in the same gene in different sugar-consuming lineages occurs more frequently than expected, we performed the equivalent analyses on matched non-sugar controls - swifts, falcons, lyre- and scrubbirds, and control Passerida lineages (Fig. 2A, fig. S3B). This control screen revealed fewer genes (404) selected specifically in these lineages, and lower proportions under selection in two (25 genes) and three (two genes) lineages (Fig. 2A). No gene shows a signal of selection in all four control groups. Using a two-sided Fisher’s exact test, we found that sugar-consuming clades show significantly more repeated selection signals than controls (Fig. 2B). To identify convergent amino acid substitutions that may not be detected by branch-level tests of positive selection, we applied CSUBST (*36*) to all 11,688 codon alignments used in the selection screen, and identified convergent amino acid substitutions among pairs of sugar-consuming clades in only 50 genes, with no gene exhibiting convergent amino acid substitutions shared by three or more sugar-consuming clades. None of 66 genes with repeated positive selection exhibited convergent amino acid substitutions. We obtained similar results for the core control groups, although we found more genes with convergent amino acid changes (232 genes with CSUBST signal; 0 overlap the 27 repeatedly positively selected genes). However, the signals in sugar-consuming lineages appear more robust, as they maintain a significantly higher normalized rate of convergent amino acid substitution (median omegaC 10.05 vs 5.79 in controls; two-sided Mann-Whitney U test, W = 74,522, P = 3.05e-13) despite having fewer overall substitutions.

**Fig. 2.**
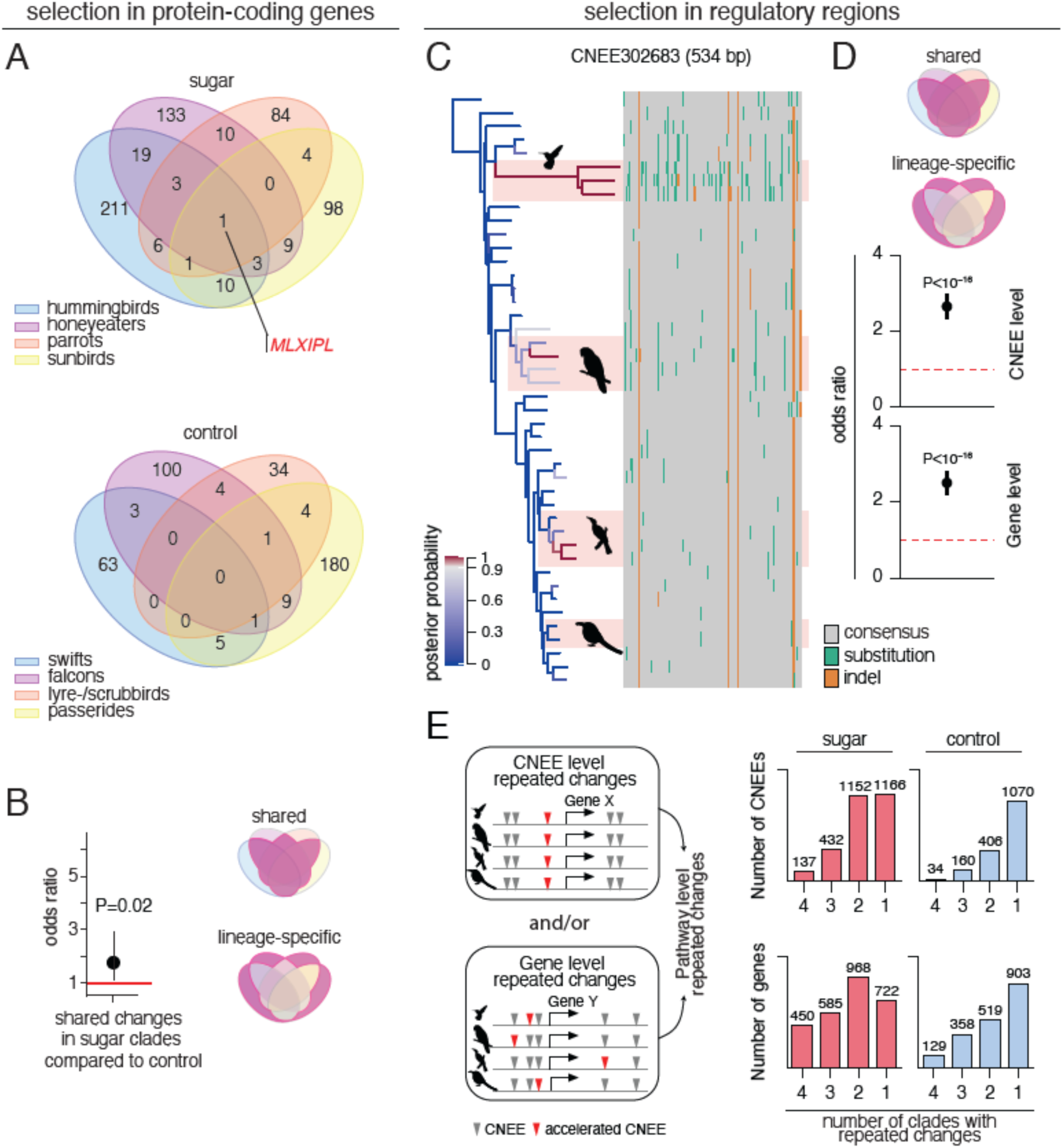
Genome-wide screens reveal higher genetic repeatability in sugar-consuming lineages in protein-coding and putative regulatory regions. (A) Number and overlap of genes under positive selection in aBSREL (on at least one branch per clade; branch-corrected p-value<0.01; no selection signal in controls) in four sugar-consuming clades and four core non-sugar-consuming control clades. (B) Excess of repeated positive selection in protein-coding genes in sugar-feeders. Counts of genes showing shared vs. lineage-specific positive selection were compared between sugar-consuming groups and matching core non-sugar controls. (C) Example of a CNEE (associated with *MAT2B*) with accelerated substitution rates in hummingbird, parrot, and honeyeater lineages. Branches in the tree (left) are colored according to the posterior probability of being in an accelerated state; those with a posterior probability >0.9 are depicted in red. Branch lengths correspond to the posterior mean of the substitution rate. The sequence alignment representation uses gray color for positions that are identical to the predominant nucleotide; turquoise and orange positions indicate deviations from the consensus (substitutions or indels). (D) Excess of repeated acceleration in sugar-feeders. Counts showing shared vs. lineage-specific acceleration were compared between sugar-consuming groups and matching core non-sugar controls for CNEEs (top) and genes associated with accelerated CNEEs (bottom), considering CNEEs accelerated in at least two clades. Significant differences are shown in black. A two-sided Fisher’s exact test was used in (B) and (D). (E) Repeated accelerations can occur at the CNEE, gene, or pathway level (left schematic). Bar plots (right) show the extent of overlap across sugar-consuming (red-filled) and core non-sugar (blue-filled) clades for accelerated CNEEs (top right) and their associated genes (bottom right).

We observed a similar excess of repeated patterns of accelerated sequence evolution in CNEEs in sugar-consuming lineages compared to control lineages. Using PhyloAcc, we identified 2,887 (0.79% of 363,476) CNEEs accelerated in at least one sugar-feeder branch: 2,014 in hummingbirds, 1,748 in parrots, 898 in honeyeaters, and 654 in sunbirds (Fig. 2C, fig. S4A). Among these, 1,152 CNEEs (40%) were accelerated in two sugar-consuming clades, 432 (15%) in three, and 137 (4.7%) across all four (Fig. 2E). Repeating this analysis for the same core non-sugar control lineages, we find fewer accelerated CNEEs and a lower proportion of repeated acceleration: 1,670 in total, with 891 in swifts; 601 in falcons; 445 in lyre- and scrubbirds; and 561 in control Passerida lineages (fig. S4B), with 406 (24%) CNEEs repeatedly accelerated in two clades, 160 (9.6%) in three, and 34 (2.0%) in all four (Fig. 2E). Two-sided Fisher’s exact tests revealed significantly more repeated CNEE acceleration in sugar-consuming clades than in controls, similar to the pattern we observe for protein-coding gene evolution (Fig. 2D, fig. S4E).

Repeated regulatory changes can also occur at the gene level, if independent clades have distinct accelerated CNEEs that are associated with the same target gene (Fig. 2E). To investigate this, we first assigned CNEEs to potential target genes, considering both a promoter-proximal and an up to 1 Mb extended domain (*42*). The 2,866 CNEEs accelerated on at least one sugar-feeder branch are associated with 2,725 genes (2,120 genes for hummingbirds; 1,959 for parrots, 1,179 for honeyeaters; and 955 for sunbirds; fig. S4D), whereas the 1,669 CNEEs accelerated in at least one core non-sugar group are associated with 1,909 genes (1,152 genes in swifts; 872 in falcons; 622 in lyre- and scrubbirds; and 885 in control Passerida species). Individual genes are significantly more likely to have associated accelerated CNEEs in multiple sugar-consuming clades than in controls. We found that 968 genes (36% of all genes with at least one accelerated CNEE) have accelerated CNEEs in two sugar-consuming clades, vs. 519 genes (27%) associated with accelerated CNEEs in two control clades, 585 (21%) vs. 358 (19%) in three clades, and 450 (17%) vs. 129 (6.8%) in all four clades (Fig. 2E, fig. S4E). Consistent with our analyses of positive selection on protein-coding genes and acceleration of individual CNEEs, a two-sided Fisher’s exact test revealed a significantly higher proportion of genes have accelerated CNEEs in multiple sugar-consuming lineages than in multiple controls lineages (Fig. 2D, 2E, fig. S4E).

## Both sequence and regulatory evolution affect carbohydrate metabolism in sugar-feeders

### Repeated and lineage-specific changes in glucose-utilization pathways in sugar-feeders

We identified only a single gene with strong evidence for positive selection in members of all four sugar-feeding lineages: *MLXIPL* (Fig. 2A, table S2). This gene encodes the transcription factor ChREBP (carbohydrate-responsive element-binding protein), which regulates cellular responses to high concentrations of glucose and fructose (*43*, *44*). Although no other genes have signatures of adaptive protein evolution in all four sugar-consuming clades, we found repeated signals of positive selection, CNEE acceleration, amino acid substitutions, and gene expression shifts, specific to two or more sugar-consuming lineages, in genes with functions in sugar intestinal absorption, cellular uptake, and cellular metabolism of sugar, all key components of glucose-utilization pathways.

Efficient sugar metabolism begins with intestinal absorption and continues through systemic transport and cellular processing. In the intestine, sucrase-isomaltase (SI) cleaves dietary sucrose into glucose and fructose, which are then imported across the apical membrane by SGLT1 (*SLC5A1*) for glucose and GLUT5 (*SLC2A5*) for fructose, followed by basolateral GLUT2 (*SLC2A2*) transporting both monosaccharides into the hepatic portal vein (*7*, *8*). We found accelerated CNEEs near *SI* in hummingbirds and honeyeaters, and duodenum-specific expression in all tested species (fig. S7), although we only detected significant upregulation relative to non-sugar-feeding controls in hummingbirds. We find a similar pattern of expression change in SGLT1: consistent duodenum-specific expression across species, paired with strong upregulation in the duodenum of hummingbirds, but downregulation in both honeyeaters and lorikeets (fig. S8). These patterns are consistent with the previous work showing that hummingbirds have the highest sucrose digestion capacity among sugar-feeders, followed by honeyeaters and sunbirds (*18*, *45*). The main fructose transporter, GLUT5, also shows high duodenal expression in all three sugar specialists, and is significantly upregulated in the duodenum of both hummingbirds and honeyeaters (Fig. 3A). While we did not detect expression changes in lorikeet duodenum, we did detect positive selection in lorikeet GLUT5 protein (Fig. 3A), suggesting GLUT5 is a repeated target of protein and regulatory evolution across all sugar-feeding clades.

**Fig. 3.**
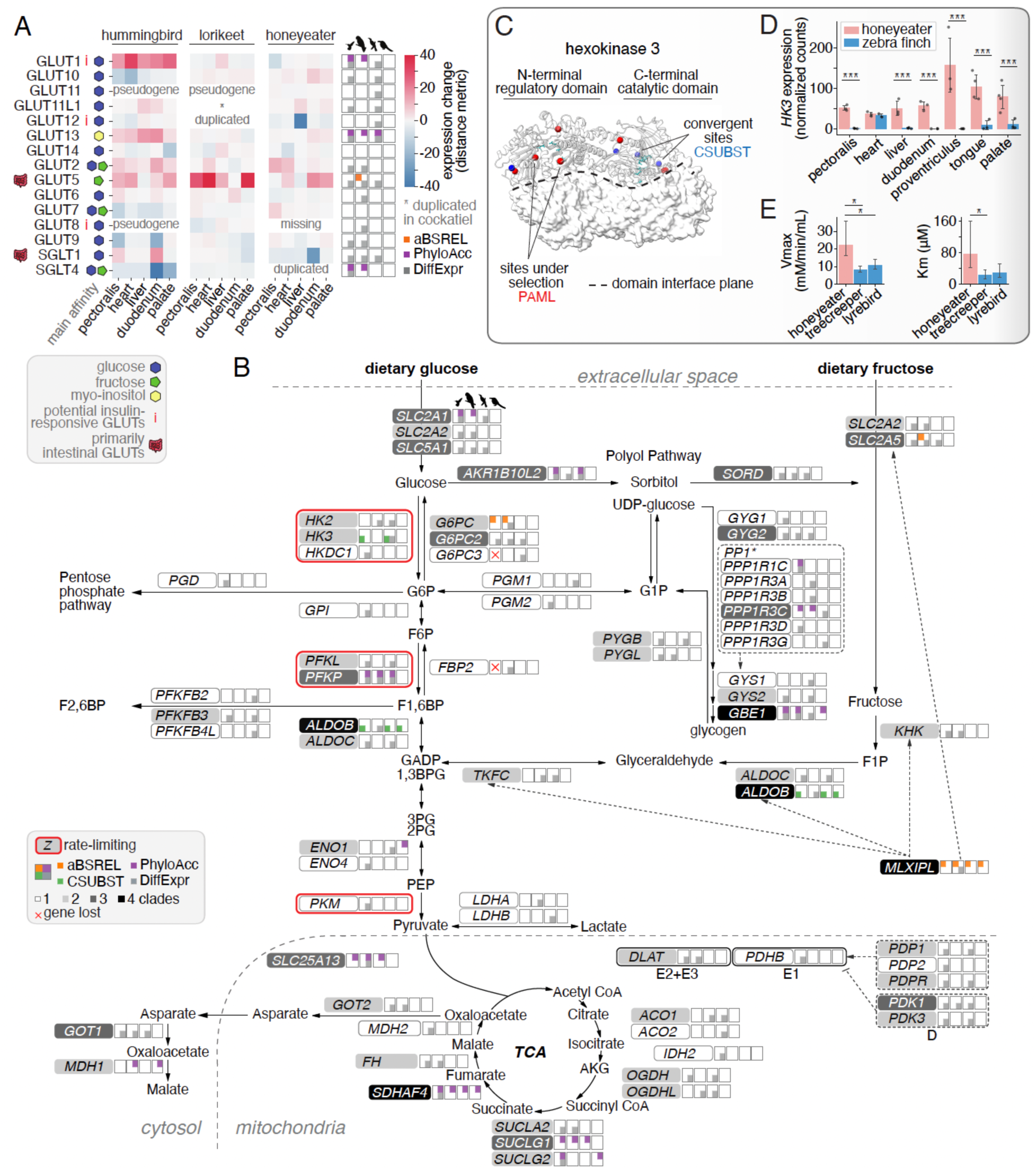
Genes involved in carbohydrate metabolism and cardiac function exhibit repeated changes across sugar-consuming clades. (A) Integrated results from aBSREL, PhyloAcc, CSUBST, and differential expression analysis for glucose transporters (GLUTs). Heatmaps show expression shifts in each sugar-consuming species compared to respective control species across all tested tissues. DiffExpr shows only top 5% differentially expressed genes, in any of the tissues. (B) Integrating results from aBSREL, PhyloAcc, CSUBST, and differential expression analysis (DiffExpr, both between-species pairwise comparisons across all analyzed tissues and multifactor cross-tissue analyses) highlights changes in sugar-feeders in glucose and fructose metabolism (glycolysis, gluconeogenesis, glycogen synthesis, TCA cycle). Signals from the four analyses are represented in the four quadrants of squares to the right of each gene name. Each square represents a sugar-consuming clade, arranged in the order of hummingbirds, parrots, honeyeaters, and sunbirds, and indicates genes under selection in at least one branch of that clade. Genes with signals from at least one of the analyses detected in all four sugar-consuming clades are shaded black, those in three clades are dark gray, and those in two clades are light gray. The rate-limiting steps of glycolysis (*50*) are highlighted with red boxes. The dotted arrows indicate some of the downstream targets of MLXIPL. Pathway schematic modified from (*101*, *106*, *107*). (C) Protein model of the honeyeater hexokinase 3 based on the human *HK3* ortholog. Sites identified as under selection by PAML and convergent amino acid substitutions identified by CSUBST are highlighted. (D) Expression of HK3 across tissues in the New Holland honeyeater compared to a non-sugar outgroup (zebra finch). *** signifies p-value < 0.001. (E) Kinetic characteristics of the honeyeater recombinant HK3 compared to control outgroups (treecreeper and lyrebird). Error bars indicate the 2.5% and 97.5% confidence intervals. Statistical significance (*) was determined by non-overlapping 95% confidence intervals.

At the systemic level, sugar uptake is facilitated by GLUT1 (*SLC2A1*), a transporter responsive to dietary status in hummingbirds (*11*) and one of the potential insulin-responsive transporters in birds (*7*, *46–48*). GLUT1 shows broad tissue expression in all tested control and sugar-consuming species, is dramatically upregulated across all tested hummingbird tissues, and harbors nearby accelerated CNEEs in both hummingbirds and lorikeets (fig. S8, Fig. 3A). The absence of GLUT1 upregulation in honeyeaters and lorikeets may reflect differences in the feeding status of sampled individuals. In contrast, GLUT5, which shows expression restricted to intestines and liver in controls, is broadly expressed across most analyzed tissues in all tested sugar-feeders (fig. S9), suggesting that it has been repeatedly and independently co-opted as a systemic fructose transporter in multiple sugar-specialist clades. Since GLUT5 is constitutively expressed (*11*), its widespread upregulation likely supports continuous fructose absorption across tissues, consistent with the ability of hummingbirds to fuel flight with fructose as effectively as glucose (*49*). The expression patterns of GLUT1 and GLUT5 suggest that sugar-feeding birds have evolved two complementary strategies for systemic sugar transport: GLUT1 for regulated glucose uptake (most prominent in hummingbirds), and GLUT5 for constitutive fructose uptake across all sugar specialists.

At the cellular level, sugar-feeding birds show repeated signatures of positive selection on key enzymes regulating sugar metabolism. *HK3* and *PFKP*, which catalyze rate-limiting steps of glycolysis (*50*), and *ALDOB* - a key enzyme in both fructolysis and glycolysis - exhibit repeated signals across sugar-consuming groups in multiple analyses (Fig. 3B). The final step of gluconeogenesis, catalyzed by glucose-6-phosphatase (encoded by *G6PC* and *G6PC2*) (*51*), shows protein sequence and regulatory evolution in at least two sugar-consuming groups (Fig. 3B). Glycolysis feeds into the tricarboxylic acid (TCA) cycle, where many genes exhibit signals of regulatory evolution across at least two sugar-feeding lineages (Fig. 3B), including *SDHAF4* that has associated accelerated CNEEs in all four sugar-consuming clades. *SDHAF4* functions at the critical interface between the TCA cycle and oxidative phosphorylation by participating in both succinate oxidation and electron transfer via complex II, thereby linking these fundamental energy-producing processes (*52*, *53*). In addition, succinate itself modulates blood pressure via the renin–angiotensin–aldosterone pathway (*54*) and promotes lipolysis in adipose tissues (*55*).

A key component of the polyol pathway encoded by *AKR1B10L2* shows accelerated CNEEs and differential expression in multiple sugar lineages (Fig. 3B). *AKR1B10L2* encodes an aldose reductase, an enzyme that facilitates the conversion of excess glucose to fructose through the intermediate sorbitol (*56*). This pathway may provide a mechanism for redirecting excess glucose into fructose metabolism, potentially enhancing the overall capacity of sugar specialists to process large quantities of dietary sugars.

### Functional shift in honeyeater hexokinase 3

CSUBST identified convergent amino acid substitutions shared between the ancestral honeyeater and the ancestral hummingbird branches in *HK3* (hexokinase 3). As a hexokinase enzyme, the HK3 homodimer catalyzes the phosphorylation of glucose in the first (and rate-limiting) step of glycolysis (Fig. 3E). While our strict positive selection screen that uses weighted p-values across all models did not show a signal of selection, the default model detected strong and specific positive selection signal in the ancestral honeyeater branch (fig. S9), and three convergently changed codons detected by CSUBST are located in close proximity to the sites identified as targets of positive selection by PAML (Fig. 3C). In addition, our expression data showed a substantial upregulation of *HK3* across all tested tissues (except heart) in honeyeaters specifically (Fig. 3D, fig. S7F). Together, this evidence suggests potential functional changes in honeyeater hexokinase 3.

Hexokinase 3 contains a catalytic C-domain and a regulatory N-domain, which face each other in the asymmetric homodimer and form two substrate-binding sites (*57*, *58*). To investigate the structural location of amino acid residues affected in the honeyeater protein, we modeled the structure of the honeyeater enzyme based on the available structure of a human hexokinase 2 as a template. Most positively selected sites and convergent amino acid substitutions are located on the interface between the catalytic and regulatory domains (Fig. 3C). Such proximity suggests that selection in the honeyeater HK3 targeted allosteric regulation and did not affect the catalytic site directly.

To assess functional changes in honeyeater HK3, we synthesized and purified the HK3 proteins from the New Holland honeyeater and two closely related outgroup species (superb lyrebird and rufous treecreeper), and conducted a hexokinase activity assay. Consistent with previous studies on HK3 from several species (*59–63*), we observed substrate inhibition, according to which enzymatic activity decreases with excess substrate after reaching a maximum turnover rate at a glucose concentration of 0.5 mM (*64*). Therefore, we compared enzymatic kinetic properties by determining Vmax (maximum velocity), Km (Michaelis constant, indicating the affinity for the substrate), and Ksi (substrate inhibition constant, indicating the concentration at which substrate begins to inhibit the enzyme). The honeyeater HK3 exhibited significantly higher Vmax (21.1 mM/min/mL) compared to control species (treecreeper: 8.0; lyrebird: 10.6 mM/min/mL; non-overlapping 95% CIs). The honeyeater HK3 also had the highest Km (65.9 μM), significantly different from the treecreeper (26.2 μM) and higher than the lyrebird (30.6 μM) (Fig. 3E, fig. S10). Honeyeater hexokinase 3 has a low affinity for glucose (high Km) and a high maximum rate, similar to mammalian glucokinase, which may make the enzyme well suited for processing glucose at high blood glucose concentrations (*65*), and may possibly reflect an adaptation to the high-sugar diet of honeyeaters.

## Shared regulatory evolution in blood pressure regulation and type 2 diabetes associated genes

CNEEs accelerated in multiple sugar-consuming lineages are significantly associated with genes that regulate blood pressure (Fig. 4A, fig. S6A-C, tables S6, S7). Given that nectar-feeding birds face the dual physiological challenge of processing high sugar concentrations while managing substantial fluid loads from their diet, this enrichment suggests coordinated regulatory evolution targeting cardiovascular homeostasis.

**Fig. 4.**
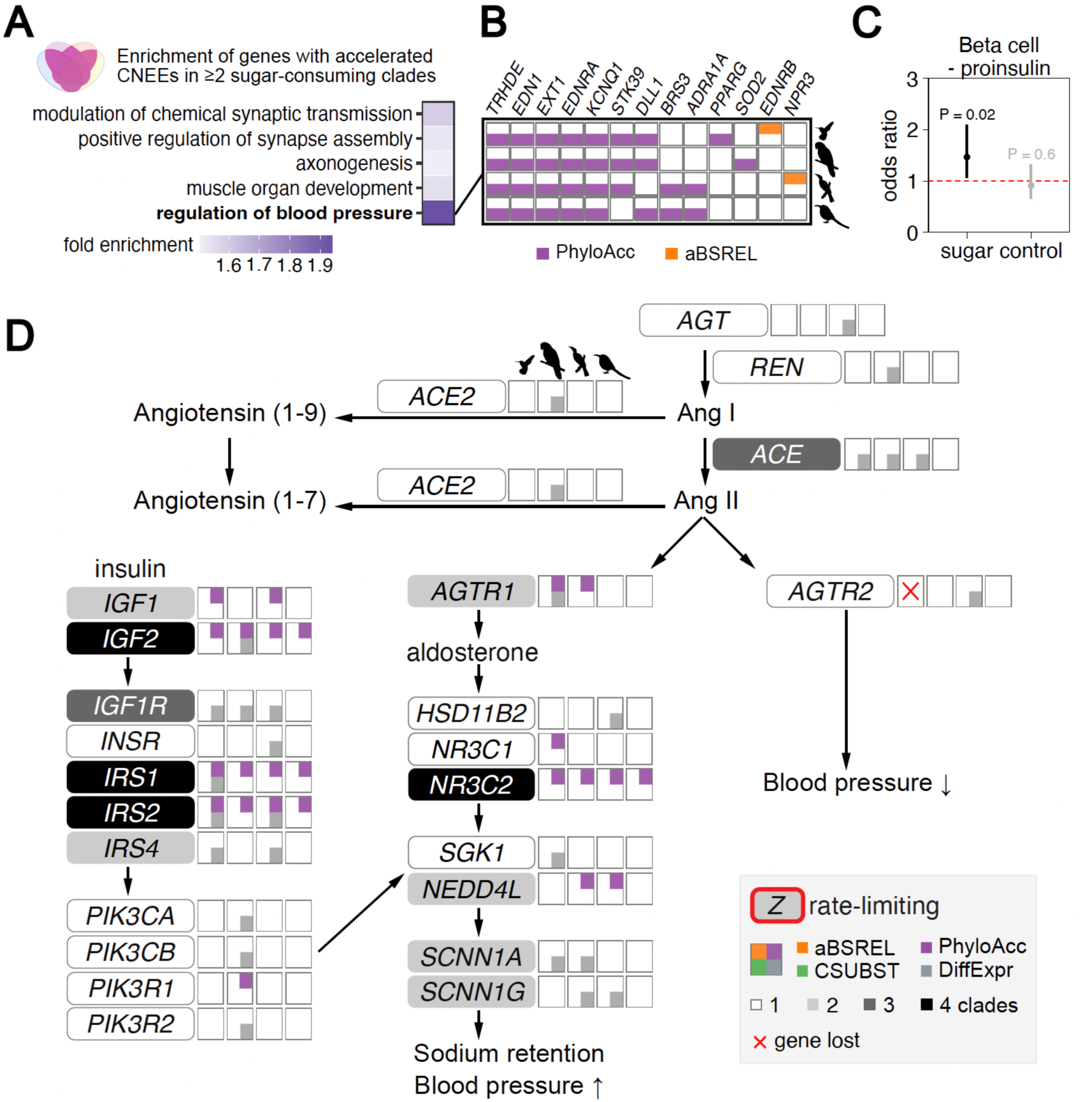
Summary of integrated results from different methods in the renin-angiotensin-aldosterone system (RAAS), insulin signaling, and type 2 diabetes associated genes. (A) GO term enrichment analysis for genes associated with accelerated CNEEs in at least two sugar-consuming clades. Only GO terms with significant results from hypergeometric tests (FDR-adjusted p < 0.3) are shown. Despite modest p-values due to small gene numbers, regulation of blood pressure shows nearly two-fold enrichment (1.94; heatmap; see also fig. S6A–B). See Table S7 for full enrichment statistics. (B) All genes annotated with the GO term regulation of blood pressure that show evidence of accelerated CNEEs (purple; PhyloAcc) or positive selection in protein-coding regions (orange; aBSREL) in sugar-consuming birds. While only two genes show positive selection, the larger number with regulatory acceleration suggests repeated modification primarily at the regulatory level. (C) Two-sided Fisher’s exact tests show enrichments of CNEEs accelerated in sugar-feeding lineages near genes associated with T2D in the cluster linked to beta cell deficiency with reduced (-) proinsulin secretion. Significance is indicated in black. (D) Integrating results from aBSREL, PhyloAcc, CSUBST, and differential expression analyses (DiffExpr) highlight changes in genes involved in the renin-angiotensin-aldosterone system, a key regulator of blood pressure and fluid homeostasis. Schematics as in Fig. 3B. Pathway modified from (*72*, *108*). The figure includes PI3K subunits that have been demonstrated to be insulin-inducible in chickens (*76*).

Key blood pressure regulating genes associated with repeated CNEE acceleration include *EDN1* (endothelin-1), which encodes a potent vasoconstrictor, and *EDNRA* (endothelin receptor A), which mediates endothelin-1 signaling in vascular smooth muscle (*66*). *KCNQ1*, encoding a cardiac potassium channel that regulates heart rhythm, also showed repeated CNEE acceleration. *KCNQ1* serves as an integrative component, as it not only controls cardiac function but also regulates ion transport in kidney and intestinal tissues, linking cardiovascular performance with electrolyte balance (*67–70*) — a particularly important function for birds managing high-volume, high-sugar diets.

Because the renin-angiotensin-aldosterone system (RAAS) is the primary mechanism coordinating blood pressure with water balance in vertebrates including birds (*71–73*), we examined RAAS components in sugar-consuming birds. We found repeated CNEE acceleration near *NR3C2* (mineralocorticoid receptor) across all sugar-consuming clades and near *NR3C1* (glucocorticoid receptor), which also binds aldosterone but with lower affinity than *NR3C2*, in hummingbirds (Fig. 4D). *AGTR1* (angiotensin II receptor type 1) showed acceleration in hummingbird and parrot clades, while *NEDD4L* (an ENaC inhibitor) was accelerated in parrot and honeyeater clades. These modifications in *AGTR1*, which mediates aldosterone signaling, and *NEDD4L*, which regulates sodium channel stability, suggest fine-tuning of sodium homeostasis mechanisms. We found that *AGTR2* is absent in all hummingbird genomes examined, a receptor that typically counteracts *AGTR1’s* effects by promoting vasodilation and natriuresis (*74*), potentially altering the balance of angiotensin II signaling.

RAAS is intricately linked with insulin signaling, a connection well established in mammals (*71*, *74*). We found that insulin/IGF-related genes showed coordinated regulatory changes. CNEEs near *IRS1, IRS2,* and *IGF2* exhibited repeated acceleration across all sugar-consuming clades, whereas CNEEs near *IGF1* additionally showed acceleration in hummingbird and honeyeater clades (Fig. 4D). Although birds exhibit unique aspects of insulin physiology, including relative insulin insensitivity compared to mammals (*75–77*), the coordinated regulatory changes we observed in RAAS and insulin/IGF signaling pathways suggest that sugar-consuming species have fine-tuned this crosstalk at the regulatory level to manage the dual challenges of high sugar and fluid loads from nectar consumption.

The evolutionary changes we observed in both metabolic and blood pressure regulation pathways led us to examine avian orthologs of human genes associated with type 2 diabetes (T2D), since T2D represents a dysregulation of glucose homeostasis and blood pressure in humans. We used data from a recent large-scale meta-analysis that identified 611 genes associated with T2D; these were grouped into eight clusters based on their genetic contributions, as measured by effect sizes of variants identified through GWAS (*78*). We tested whether genes dysregulated in T2D in humans overlap genes likely involved in sugar diet adaptations in birds. Indeed, we found a significant overlap of CNEEs showing shared accelerations in sugar-feeding birds with T2D associated genes in the dysfunctional beta cell cluster, while CNEEs accelerated in non-sugar-feeders showed no such enrichments (two-sided Fisher’s exact test P=0.02, Fig. 4C). Overall, 34 genes associated with T2D in humans are also associated with CNEEs that are independently accelerated in all four sugar-consuming clades. These include a number of genes that are crucial mediators of insulin signaling, beta cell function, and glucose homeostasis such as *IRS1, IRS2*, *IGF2*, and *KCNQ1* (table S8) (*79–82*). However, we did not find evidence for repeated positive selection across multiple independent sugar-consuming clades in any genes associated with T2D in humans. These findings reinforce the idea that evolutionary responses to a sugar-rich diet in birds substantially involve regulatory modification of genes affecting insulin secretion and cardiovascular homeostasis.

## Large effects of regulatory evolution in lipid and amino acid metabolism

To investigate expression changes associated with a high-sugar diet, we conducted two complementary transcriptomic analyses: (i) a between-species analysis comparing pairs of sugar-feeding birds and their corresponding controls, and (ii) a multifactor cross-tissue analysis to detect tissue-specific expression shifts (Fig. 1F). Genes with upregulated expression in sugar-feeders show enrichments in many processes associated with energy derivation across all tested tissues (table S11), with the term organic acid biosynthesis and pyruvate metabolism repeatedly enriched in at least two sugar-consuming clades in the liver (Fig. 5A, table S11). This is also supported with our cross-tissue analysis where sugar-specific upregulations in liver and pectoralis are repeatedly enriched in mitochondrial function and metabolic energy generation terms which encompasses glycolysis, electron transport chain, and other pathways involved in energy generation in cells (Fig. 5C).

**Fig. 5.**
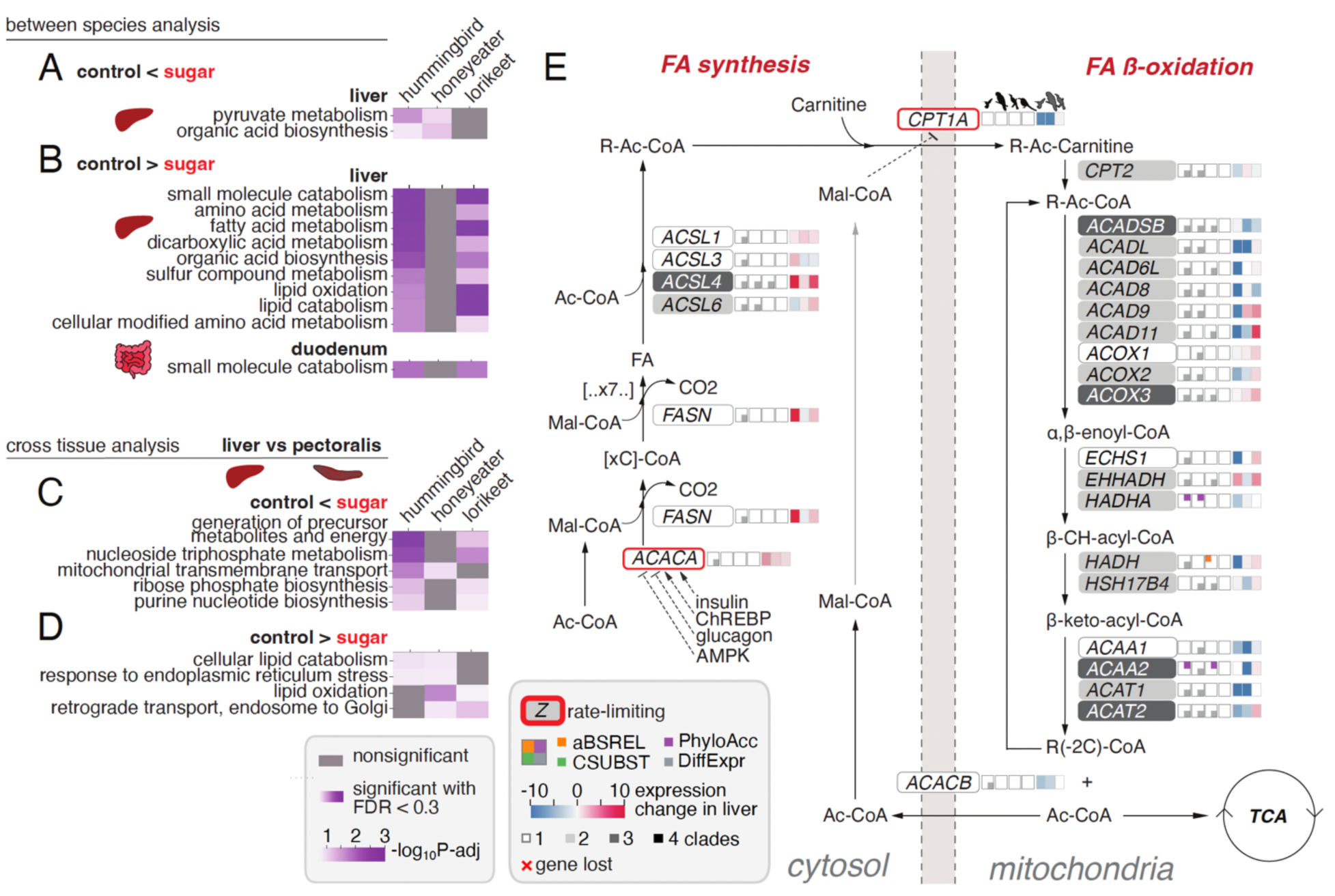
Sugar-feeders show large changes in the regulation of lipid and amino acid metabolism. (A, B) GO Biological Processes significantly enriched (FDR<0.3) for (A) up- and (B) downregulated genes in at least two sugar-feeders in the pairwise differential expression analysis between species. (C, D) GO Biological Processes significantly enriched (FDR<0.3) for genes with tissues-specific expression that show (C) up- and (D) downregulation in sugar clades compared to controls. Only pathways showing enrichment in at least two sugar clades are shown. (E) Large effects of regulatory evolution in lipid metabolic pathways and contrasting expression signals in fatty acid synthesis and beta-oxidation. Schematics as in Fig. 3D. The heatmap panels to the right of the quadrants illustrate the differential expression in the liver of hummingbirds, parrots, and honeyeaters compared to their respective outgroups. Pathway modified from (*109*, *110*).

Among downregulated genes, we observed a consistent enrichment of genes involved in lipid metabolic processes for liver, duodenum, and pectoralis in all sugar-feeders (Fig. 5B, D, table S12). This expression shift is substantially driven by the downregulation in multiple sugar-consuming species of genes with roles in fatty acid metabolism, including *ASPG* (hummingbird, lorikeet), *DECR2* (hummingbird, lorikeet, honeyeater), *ECHDC1* (hummingbird, lorikeet), *FABP1* (hummingbird, lorikeet, honeyeater), and *HACL1* (hummingbird) (table S10). Additionally, genes downregulated in the liver are enriched in amino acid metabolism in at least two sugar-feeding groups (Fig. 5B, table S12). Many genes involved in amino acid metabolism belong to the branched chain amino acid (BCAA) pathway; these include: *BCKDHB* (hummingbird, honeyeater), *ASRGL1* (hummingbird, honeyeater, lorikeet), *MCCC2* (hummingbird, lorikeet), *GOT1* (hummingbird, honeyeater), *GOT2* (hummingbird, lorikeet), and *ALDH4A1* (hummingbird, lorikeet) (fig. S11).

Examining genes involved in lipid metabolism in more detail, we found that most genes involved in fatty acid (FA) synthesis and FA beta-oxidation showed contrasting signatures of regulatory evolution in at least one sugar-consuming lineage (Fig. 5E). In particular, we observed a striking upregulation of genes responsible for FA synthesis in the liver of all sugar-consuming clades, while genes responsible for FA beta-oxidation are downregulated. For example, *ACACA* (acetyl-CoA carboxylase), encoding the enzyme that catalyzes the rate-limiting step in FA synthesis, shows repeated upregulation in all sugar-consuming clades (fig. S7E).

Highlighting the expression shifts in amino-acid and lipid metabolic pathways, we also found significant downregulation in one or more sugar-consuming species for a number of genes with roles in both BCAA and lipid metabolism, including *ACADSB* (hummingbird, lorikeet, honeyeaters), *ACAD6L* (honeyeater, hummingbird), *ACAD8* (hummingbird), *ACAT1* (hummingbird, lorikeet), *PCCA* (hummingbird, lorikeet), and *PCCB* (hummingbird, lorikeet) (fig. S11). Because flower nectar is poor in protein and fat, sugar-feeders may have to conserve these nutrients. These results suggest that sugar-feeders impose breaks on amino acid and lipid catabolism across all tissues. In addition, higher levels of free amino acids in sugar-feeding birds may protect them from protein glycation which is a major risk associated with high sugar intake (*83*). The observed repeated downregulation of genes involved in amino acid degradation across sugar-feeding clades suggests a potential mechanism to preserve amino acids and mitigate glycation-related damage.

One of the genes involved in lipid metabolism regulation showing the strongest signal was identified in the context of systemic transporters. *GLUT13* (SLC2A13) is strongly upregulated across all hummingbird tissues and has accelerated CNEEs near its locus in hummingbirds, honeyeaters, and parrots (Fig. 3A). *GLUT13* transports myo-inositol, a signaling molecule shown to regulate fat metabolism in mammals (*84*).

## Functional changes in sugar-feeders’ *MLXIPL* enhance sugar response

There is exactly one gene that is robustly positively-selected in members of all four sugar-consuming clades and no non-sugar control lineage: *MLXIPL*. This gene encodes the transcription factor ChREBP that is activated by glucose metabolites and dimerizes with its interaction partner MLX to bind to DNA, initiating the expression of target genes that promote glucose metabolism and lipogenesis, thus enhancing the capacity to metabolize ample amounts of sugar (*85*, *86*) (Fig. 6A). *MLXIPL* as well as downstream target genes are involved in de novo lipogenesis, glycogen synthesis, and glucose and fructose metabolism (Fig. 6B and fig. S12B), and are upregulated in all three sugar-consuming clades relative to non-sugar outgroups. In hummingbirds, *MLXIPL* expression in the duodenum was the highest among all tissues analyzed (Fig. 6C), possibly indicating the greater importance of intestinal compared to hepatic *MLXIPL* expression for fructose tolerance (*87–90*). Hummingbirds also show exceptionally high hepatic expression of lipogenic target genes of ChREBP, which are associated with a dramatic accumulation of lipids in their livers (Fig. 6D). We observed that many amino acid substitutions specific to sugar-consuming clades cluster in the putative glucose-response activation conserved element (GRACE domain) (*91*), which is thought to mediate transcriptional activity in response to glucose (*92*) (Fig. 6E).

**Fig. 6.**
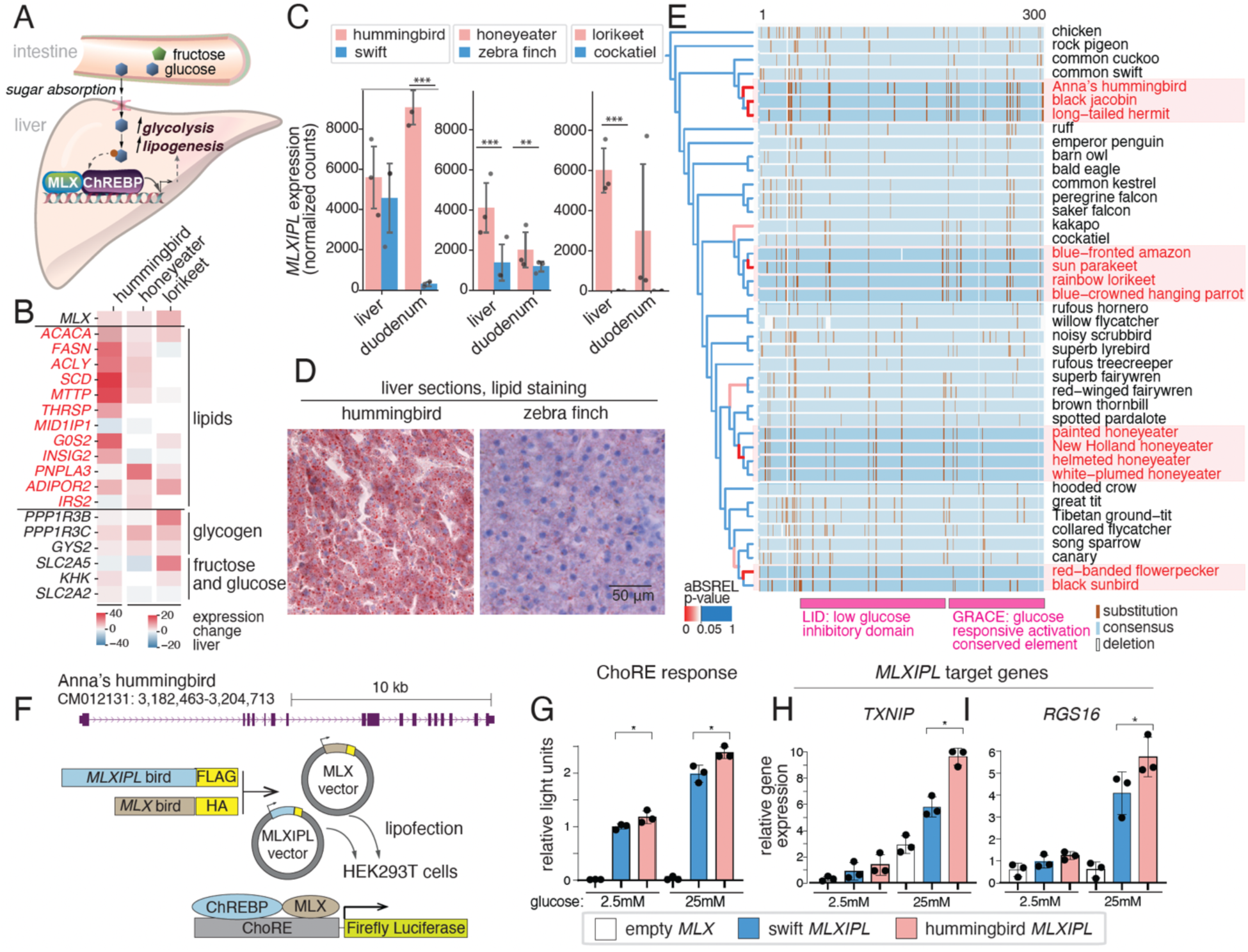
Positive selection and functional changes of MLXIPL. (A) Role of MLXIPL-encoded carbohydrate response binding protein (ChREBP) in the intestine and liver. (B) Expression changes in MLXIPL target genes involved in lipid and sugar metabolism in liver of sugar-consuming species compared to their corresponding non-sugar-consuming controls. (C) *MLXIPL* expression in liver and duodenum of sugar-consuming species compared to their corresponding non-sugar-consuming controls. (D) Oil-red-O lipid staining of liver sections. Red droplets represent lipid deposition. Nuclei and cytoplasm are counterstained with hematoxilin (purple). (E) Part of MLXIPL protein alignment overlapping known glucose-sensing domains. Substitutions are marked with brown lines. Branches in the phylogeny are colored according to the aBSREL branch-corrected p-value (red branches are p-value < 0.05). (F) Vector design and reporter assay to assess transcriptional activity of the recombinant MLXIPL proteins. (G) Transcriptional activity of the long MLXIPL isoforms in the luciferase reporter assay with the hummingbird and swift MLXIPL-MLX recombinant protein pairs. (H, I) Expression changes in two target genes of MLXIPL. In (C) and (G-I) * signifies p-value < 0.05, ** - p-value < 0.01, and *** - p-value < 0.001

To test the functional consequences of these sequence changes we detected in sugar-feeders, we synthesized hummingbird and swift *MLXIPL* together with its corresponding interaction partner *MLX*. We expressed these vectors ectopically in human embryonic kidney cells (HEK293T) (Fig. 6F) and used a carbohydrate response element (ChoRE)-driven luciferase reporter assay (*93*) to test two predominantly expressed *MLXIPL* isoforms (fig. S12C,D). We found that for both birds, the long isoform drives a much higher ChoRE response compared to the short one, suggesting the long isoform is the main functional isoform in birds (Fig. 6G, fig. S12E,I).

We found that hummingbird ChREBP shows a higher ChoRE response compared to that of the swift at both low and high concentrations of glucose (Fig. 6G, fig. S12I), a difference that was also reflected in the higher capacity of the hummingbird protein to drive expression of the ChREBP target genes *TXNIP* (*94*) and *RGS16* (*95*) (Fig. 6H,I, fig. S12J,K). This suggests that a given glucose concentration will drive a higher expression of ChREBP target genes in hummingbirds and thereby increase their cellular capacity to tolerate and metabolize simple carbohydrates. Hence, both expression levels (Fig. 6C) and transcriptional activity of the MLXIPL protein (Fig. 6G) may enable hummingbirds to specialize on high-sugar diets.

## Discussion

Using multiple independent transitions to sugar-rich diets as a system, we addressed two fundamental questions: (i) whether repeated molecular changes, lineage-specific changes or both contribute to convergent adaptations and (ii) what are genomic adaptations to sugar-rich diets in birds. Our comprehensive analyses integrating genomic screens for multiple evolutionary signatures revealed that both repeated *and* lineage-specific molecular changes contribute to adaptations to these specialized diets. Furthermore, these adaptations involve repeated evolution in both regulation (detected with PhyloAcc and DESeq2) and proteins (detected with aBSREL and CSUBST). Specifically, in comparison to non-sugar-consuming control lineages, we found significantly more repeated positive gene selection and repeated accelerated evolution in putative regulatory regions (CNEEs) in multiple sugar-consuming clades. These repeated genomic changes show considerable functional coherence, and are associated primarily with energy homeostasis, metabolism, blood pressure regulation, and insulin signaling, pointing to a common molecular toolkit required to adapt to high-sugar diets.

Our comprehensive genome-wide analyses of both protein-coding and regulatory evolution revealed differential contributions to different relevant biological processes. Whereas repeated protein sequence evolution in sugar-feeding birds is largely confined to genes involved in sugar metabolism, repeated regulatory evolution has occurred more broadly, targeting pathways related to blood pressure regulation, lipid and amino acid metabolism, and genes associated with type 2 diabetes. This is consistent with previous work highlighting the importance of repeated evolution at regulatory loci for the evolution of convergent phenotypes (*39*, *96–98*). However, it is important to consider that genomic screens are imperfect and comparisons between regulatory and protein evolution in particular require caveats. Our screen for protein-coding selection was intentionally conservative, to mitigate high false-positive rates reported in previous studies (*99*, *100*), but this may lead to missing weak signals. We also recognize that some of the expression changes we observe may represent a latent, plastic response to sugar consumption, potentially accessible to non-sugar-specialist species when exposed to a high-sugar diet. However, we think this is unlikely to fully explain the signal we observe for two related reasons. First, many of the expression changes we observe are downstream of *MLXIPL*, which is positively selected in all four sugar-consuming clades, and which we confirm is functionally diverged in hummingbirds. Second, for many genes involved in glycolysis and glucose metabolism, in particular, we see evidence for concurrent acceleration of nearby CNEEs and expression change, suggesting sequence evolution and not a purely plastic response underlying these changes. Nonetheless, we acknowledge that distinguishing plastic responses from evolved responses remains an important challenge for comparative work.

In some cases, our analysis identifies repeated signals of evolution in genes previously highlighted in only a single clade. For example, our study revealed that protein sequence and expression variation of the main fructose transporter GLUT5 and key genes of the fatty acid biosynthesis are not confined to hummingbirds (*11*, *24*, *25*), but are also found in honeyeaters, sunbirds, and sugar-feeding parrots, suggesting that evolution of GLUT5 may be a widely important adaptation to a high sugar diet. Perhaps even more strikingly, we identify repeated evolutionary changes in sugar-consuming bird lineages in genes previously identified as evolving in sugar-consuming bats. These include sequence and regulatory changes in a key glycolytic gene *ALDOB* (*101*, *102*), several glucose transporters (GLUT2, GLUT5, and SGLT1) (*103*) as well as regulatory changes in gluconeogenic gene *G6PC* (*102*), and highlight the possibility that similar evolutionary changes may contribute to sugar adaptation across large evolutionary distances.

We also identify examples of molecular changes that appear to be primarily relevant in a single sugar-consuming lineage, suggesting that not all evolutionary change associated with diet evolution occurs in the same set of genes. However, the lineage-specific signals we do see are often gene-level elaborations of the same pathways in which we detected repeated signals of selection. A striking example of this is the lineage-specific change in hexokinase 3, a rate-limiting glycolytic enzyme, that was targeted by both protein and regulatory evolution in honeyeaters. Our functional assays demonstrated a shift in catalytic function in the honeyeater enzyme. Analogous to “many roads lead to Rome”, the observed lineage-specific changes in the same pathways suggest that adaptations in physiological and metabolic processes can also be realized by different genes within broad functional networks.

Many of the most compelling adaptations we observe are shared among all four clades. A particularly important example is the repeated positive selection signature in *MLXIPL*, encoding the master transcription factor ChREBP that senses glucose metabolites and regulates energy homeostasis. Through functional assays, we demonstrated how sequence changes in *MLXIPL* could contribute to substantial shifts in energy homeostasis in sugar-feeders. We hypothesize that *MLXIPL* may be a critical trigger in the upregulation of lipid synthesis, as it activates the rate-limiting enzyme of the pathway (acetyl-CoA carboxylase alpha) and multiple other genes involved in lipogenesis. The observed repeated upregulation of lipogenesis and downregulation of fatty acid catabolism likely reflects the energetic and substrate preferences in avian sugar-feeders: during feeding, birds convert glucose into fat rather than heavier hydrophilic glycogen, and during fasting, they use stored fat, upregulating lipid degradation pathways (*104*, *105*).

Overall, our results suggest that adaptations to high-sugar diets in birds require modifications to a constrained set of organismal functions, including pathways directly involved in metabolism, as well as previously-undetected pathways such as blood pressure regulation to cope with the physiological consequences of high sugar consumption. More generally, our study highlights how a genome-wide integrative analysis, considering evolutionary signatures at the gene sequence, regulatory and expression level across multiple lineages, can provide comprehensive insights into repeated as well as lineage-specific genomic adaptations relevant for a convergent trait.

## Materials and Methods

### Sample collection

#### Bird specimens for genome and transcriptome sequencing

Genomes were sequenced from tissue or blood samples from 9 species (black jacobin, common swift, rainbow lorikeet, blue-crowned hanging parrot, cockatiel, New Holland honeyeater, white-plumed honeyeater, brown thornbill, and rufous hornero) representing three of the four focal sugar-consuming groups as well as outgroup lineages. Transcriptomes from species in three sugar-consuming groups (Anna’s hummingbird, rainbow lorikeet, New Holland honeyeater) and three outgroup species (common swift, cockatiel, zebra finch) were sequenced from a replicate panel of tissue samples. Details of specimens used for each target clade are described below.

#### Hummingbirds and swifts

For hummingbirds and their closest relatives, the swifts, we sequenced the genome of a black jacobin (*Florisuga fusca*) using a liver tissue sample collected in Santa Teresa, Espirito Santo, Brazil with the permission of the Brazilian Institute of Environment and Renewable Natural (IBAMA) under the Biodiversity Information and Authorization System (SISBio) permit numbers 30319-1, 41794-2 and 49097-1 (Genetic Heritage #A8E6064). A series of tissues for transcriptomic analysis was collected from three male Anna’s hummingbirds (*Calypte anna*) collected in Vancouver, British Columbia following permits obtained from Environment and Climate Change Canada under Scientific Permit SC-BC-2021-0001-01. Tissue samples from four common swifts (one individual was used for genome sequencing, three for transcriptomics) were obtained from terminally-injured individuals in the care of a rehabilitator in Germany that were euthanized at a veterinary clinic (following the regulations of the European Union and the German Animal Welfare Regulation Governing Experimental Animals (TierSchVersV) and the guidelines of the Government of Upper Bavaria).

#### Parrots

For the parrot clade, DNA was obtained from a blood sample of a captive rainbow lorikeet (*Trichoglossus moluccanus*) from the Loro Parque zoo in Tenerife and used for genome sequencing. Tissue samples for RNA-seq collected from four captive-bred rainbow lorikeet individuals from Sydney, Australia, as well as a tissue sample from a captive-bred cockatiel (*Nymphicus hollandicus*) used for genome sequencing were collected following procedures approved by the Animal Welfare Committee (AWC) at Deakin University Australia and following permits from the Department of Sustainability and Environment of Victoria, Australia; tissues from each individual are accessioned into the Museum of Comparative Zoology at Harvard University (MCZ Ornithology # 364103, 364104, 364108, 364107). An additional lorikeet genome (not part of the main analysis, but generated using the same procedure as the thornbill and New Holland honeyeater genomes (described below) and uploaded on NCBI) was generated from the individual accessioned under MCZ 364103; a tissue sample from this individual was also used for Hi-C scaffolding for the primary lorikeet genome. Cockatiel tissues for transcriptomic analysis were obtained from three individuals bred in captivity in Germany (following the regulations of the European Union and the German Animal Welfare Regulation Governing Experimental Animals (TierSchVersV)). DNA extracted from a blood sample from a blue-crowned hanging parrot (*Loriculus galgulus*) obtained from the Schoenbrunn Zoo in Vienna and kept in aviaries in Seewiesen, Germany was used for genome sequencing.

#### Honeyeaters and other passerines

For the honeyeater clade, tissue samples from two honeyeater species and a thornbill were collected following approval by the Animal Welfare Committee (AWC) at Deakin University, Australia, and according to permits from the Departments of Sustainability and Environment of Victoria and New South Wales, Australia. A tissue sample of a white-plumed honeyeater (*Ptilotula penicillata*) was collected at Fowlers, Gap, Australia (accessioned under MCZ 364075). A brown thornbill (*Acanthiza pusilla*; tissue accessioned under MCZ 364070) as well as three New Holland honeyeaters (*Phylidonyris novaehollandiae*) were collected in Geelong, Victoria: an individual with material accessioned under MCZ 364095 was used for genome and transcriptomes; two other individuals (accessioned under MCZ 364096 and MCZ 364097) were used for transcriptomes. Tissue samples from three zebra finches were collected from animals bred in captivity in Seewiesen, Germany, according to the regulations of the European Union and the German Animal Welfare Regulation Governing Experimental Animals (TierSchVersV) and used for transcriptomics. A blood sample from a wild-caught rufous hornero (*Furnarius rufus*) was collected following permits from the ethical committee of animal use (CUEA) of the Faculty of Sciences in the Universidad de la República (Nr. 186) and the Ministry of Environment of Uruguay (Nr. 138/6) and used for genome sequencing.

### Defining analysis groups and species selection for whole genome analysis

Four focal sugar-consuming groups were examined, called here ‘core sugar clades’, which are composed of species with a diet consisting of at least 50 percent nectar and fruit combined, using information from the EltonTraits database (*13*). The first clade, hummingbirds, represents a clear instance of a single origin of nectarivory, including here the early-branching lineages of jacobins and hermits (represented by the newly-sequenced black jacobin, as well as our recently-sequenced genome of a long-tailed hermit (*23*)) together with the well-studied and distantly related Anna’s hummingbird (*31*). The exact origin of nectarivory in the other clades is less clear-cut, motivating us to include species with high levels of fruit as well as nectar consumption. In parrots, multiple origins of nectarivory likely exist (including in lorikeets, *Loriculus* hanging parrots, New World parakeets, swift parrots, as well as kākās) (*13*, *14*); in addition, many species also consume large amounts of sugar-rich fruits. We focused on the rainbow lorikeet, an extreme nectarivore (like many of the closely-related lories), and included a hanging parrot representative (*Loriculus galgulus*) as well as the highly frugivorous sun parakeet and blue-crowned amazon in our focal clade. For honeyeaters, we sequenced two species of honeyeater (New Holland and white-plumed), and included the published genomes of the painted honeyeater (*30*) and the helmeted honeyeater (*111*). A genome of the brown thornbill, a member of the sister group to honeyeaters (including thornbills and pardalotes) was also sequenced; as many species in this group are largely insectivorous, but also take nectar and lerp, this lineage was classified as ‘ambiguous’. This category was also used for other species close to but outside of the primary in-group sets whose diet contained fruit or nectar (but less than 50%), and for which it was unclear if some adaptations for sugar-feeding could have already have evolved, given either their phylogenetic position and diet or other information (such as data on taste receptor function). Lastly, the published genomes of the red-banded flowerpecker and black sunbird (*30*) comprised the fourth focal clade, the ‘sunbird’ clade.

Outgroup sets (‘core non-sugar’) were selected from closely related clades where nectar and fruit are not a part of the diet: two swifts (for the hummingbird clade), three falcons (for the parrot clade), a lyrebird and scrubbird (Menuroidea; here called ‘lyre- and scrubbirds’) (the outgroup clade for honeyeaters), and zebra finch, Tibetan ground-tit, and collared flycatcher (here called ‘control Passerida’) for the sunbird clade. Lineages not classified as core sugar, non-sugar-consuming, or ambiguous, were considered as background species for the genomic analyses. Background lineages were selected from publicly available NCBI genomes to ensure balanced representation across the bird phylogeny and were chosen on the basis of having a BUSCO gene annotation completeness of at least 80%. Outgroups for transcriptomic analyses differed slightly from genomic analyses because of tissue availability: as an outgroup for rainbow lorikeets, the cockatiel, an entirely granivorous member of the non-sugar-consuming Cacatuidae family of parrots (and part of the ‘ambiguous’ set for the genomic analyses) was selected. As a note, as sugar-consuming behavior has been observed in some Nestor parrots (e.g., kākā, *N. meridionalis*), and as cacatuids as well as kakapos, kākās and keas take fruit (*13*, *14*), nectar and fruit-taking behavior may have existed in the ancestor of all parrots prior to the divergence of Cacatuidae and Psittacidae. For honeyeaters, the distantly-related granivorous zebra finch (part of the control non-sugar-consuming Passerida set for genomic analyses) was used as an outgroup.

### Tree Topology

For our analyses, we constructed an avian phylogeny based on recent phylogenomic studies. We used the phylogenomic supertree from (*112*) as the backbone topology and incorporated clade-specific relationships from specialized studies: (*113*) for hummingbirds, (*114*) for Psittaciformes, (*115*) for Meliphagidae (honeyeaters), and (*116*) for Passeriformes relationships. This topology was used consistently across all comparative analyses in our study.

### Genome sequencing

#### Acanthiza pusilla, Phylidonyris novaehollandiae

Genomic DNA was isolated from tissue samples collected as described above and accessioned in the Museum of Comparative Zoology Ornithology collection from a brown thornbill, *Acanthiza pusilla* (MCZ catalog number: 364070), and a New Holland honeyeater, *Phylidonyris novaehollandiae* (364095). DNA was isolated using the DNeasy Blood and Tissue Kit (Qiagen, Hilden, Germany) following the manufacturer’s protocol. We measured DNA concentration with a Qubit dsDNA HS Assay Kit (Invitrogen, Carlsbad, USA) and performed whole-genome library preparation, sequencing, and assembly (following (*117*, *118*)). In brief, a DNA fragment library of 220-bp insert size was prepared using the PrepX ILM 32i DNA Library Kit (Takara), and mate-pair libraries of 3-kb insert size were prepared using the Nextera Mate Pair Sample Preparation Kit (cat. No. FC-132-1001, Illumina). We performed DNA shearing for the fragment and mate-pair library preparations using Covaris S220. We used the 0.75% agarose cassette in the Pippin Prep (Sage Science) for size selection of the mate-pair library (target size 3 kb, “tight” mode). We then assessed fragment and mate-pair library qualities using the High Sensitivity D1000 ScreenTape for the Tapestation (Agilent) and High Sensitivity DNA Kit for the Bioanalyzer (Agilent), respectively, and quantified the libraries with qPCR (KAPA Library Quantification Kit) prior to sequencing. We sequenced the libraries on an Illumina HiSeq 2500 instrument (High Output 250 kit, PE 125-bp reads) at the Bauer Core facility at Harvard University.

#### Florisuga fusca

Genomic DNA (gDNA) was isolated from a female black jacobin’s liver using the MagAttract HMW DNA Kit (Qiagen, Hilden, Germany), as per manufacturer instructions. The specimen was captured and euthanized according to Monte et al., 2023 (*119*), under the capture permits 41794-1 and genetic patrimony register number AF0CC37, licensed by Ministério do Meio Ambiente, Brazil. Whole-genome shotgun libraries were prepared using Illumina’s TrueSeq 2 kit, creating paired-end libraries with 500 and 800 bp fragment sizes, as well as a 3500 bp Nextera mate pair library. Sequencing was executed on an Illumina HiSeq2500 system, and mate pair reads were processed using NEXTCLIP (*120*).

#### Furnarius rufus and Trichoglossus moluccanus

High-molecular-weight genomic DNA was extracted using the MagAttract HMW DNA Kit (Qiagen) following the manufacturer’s protocol. For *Furnarius rufus*, genomic DNA was isolated from blood preserved in Queen’s buffer, while for *Trichoglossus moluccanus*, a blood sample preserved in ethanol was used. For both species, 10X Genomics libraries were prepared using the Chromium Genome Kit (v2 chemistry) and sequenced on an Illumina NovaSeq 6000 platform (151 bp paired-end reads with 8 bp dual index). Additionally, for *T*. *moluccanus*, a Hi-C library was constructed (using a tissue sample from another lorikeet individual, accessioned into the MCZ (364103)) using the Dovetail Hi-C Kit (cat# 21004, Manual v.1.01 2-01-18). For this, 33 mg of flash-frozen powdered muscle tissue were used. The resulting barcoded library was sequenced using the same Illumina platform and configuration to ca. 50 M reads.

#### Ptilotula penicillata, Nymphicus hollandicus, Loriculus galgulus, and Apus apus

##### Extraction of ultra-long genomic DNA

High molecular weight (HMW) genomic DNA (gDNA) from muscle tissues (collected on dry ice and preserved at -80) of one common swift (*Apus apus*) and one white-plumed honeyeater (*Ptilotula penicillata*) individual were extracted following the Bionano Prep Animal Tissue DNA Isolation Fibrous Tissue Protocol (Document Number 30071, document revision C). Briefly, 30-40 mg of muscle tissue was cut into 3 mm pieces and fixed with 2% formaldehyde and Homogenization Buffer provided by the kit. The mildly fixed tissue was homogenized with a Qiagen Tissueruptor and the homogenized tissue was embedded in 2% agarose plugs. The plugs were treated further with Proteinase K and RNase A, and then washed with 1X Wash Buffer from the kit and home-made 1X TE Buffer (pH 8.0). DNA was recovered from plugs and purified finally by drop dialysis against TE Buffer (pH 8.0).

Using the Bionano SP Blood and Cell DNA Isolation Kit (PN 80030, Bionano Genomics Inc, USA), HMW gDNA from blood samples preserved in ethanol of one blue-crowned hanging parrot (*Loriculus galgulus*) was isolated applying the modified Bionano Prep SP Frozen Human Blood DNA Isolation Protocol (Document Number 30246, document revision C). In brief, the ethanol supernatant was firstly removed and the collected blood pellet was washed 3 times in 1 ml of 1 x PBS buffer followed by resuspending the pellet in cell buffer. After cell lysis with Proteinase K, the gDNA was precipitated by isopropanol and attached to the Nanobind disk. Several times of washing steps were performed and the purified gDNA was finally eluted in the Elution Buffer.

gDNA of one cockatiel (*Nymphicus hollandicus*) was extracted from muscle tissue that was preserved in ethanol using the Circulomics Nanobind Tissue Big DNA kit (Circulomics Inc., PacBio, USA) according to the manufacturer’s instructions. For initial ethanol removal, the tissue was washed three times for 30 min each in 45 ml of EtOH removal buffer (40nM NaCl, 20nM Tris (pH7.5) and 30mM EDTA).

gDNA concentration was determined using the Qubit dsDNA BR (Broad Range) assay kit (ThermoFisher, USA). The purity of the sample was measured with the NanoDrop, assessing curve shape and the 260/280 nm and 260/230 nm values. The integrity of the HMW gDNA was determined by pulse field gel electrophoresis using the Pippin PulseTM device (SAGE Science). The majority of the long and ultralong gDNA was between 50 and 500 kb in length. All pipetting steps of ultra-long and long gDNA have been done very carefully with wide-bore pipette tips.

##### PacBio continuous long read (CLR) library preparation and sequencing

For *Apus apus*, *Ptilotula penicillata*, and *Loriculus galgulus*, long insert libraries were prepared as recommended by Pacific Biosciences (PacBio) according to the ‘Guidelines for preparing size-selected >30 kb SMRTbell templates making use of the SMRTbell express Template kit 2.0. Briefly, ultra-long gDNA was sheared to 75 kb fragments with the MegaRuptor device (Diagenode) and 4-8 µg sheared gDNA was used for library preparation. The PacBio SMRTbell library was size-selected for fragments larger than 26 kb, 26kb, 20kb and 30kb with the BluePippinTM device, respectively, according to the manufacturer’s instructions. All size selected libraries were sequenced for 15 hours on the Pacbio Sequel^®^ II on 8M SMRT™ cells using the Sequel® II Binding 2.0 and the Sequel^®^ II Sequencing 2.0 chemistry.

##### PacBio High Fidelity (HiFi/CCS) library preparation and sequencing

For *Nymphicus hollandicus*, the gDNA entered library preparation using a PacBio HiFi library preparation protocol “Preparing HiFi Libraries from Low DNA Input Using SMRTbell Express Template Prep Kit 2.0”. Briefly, gDNA was sheared to 8 kb with the MegaRuptor device (Diagenode) and 2.6 µg sheared gDNA was used for library preparation. With Ampure beads size-selection, fragments shorter than 3 kb were removed. The size-selected library was prepared for loading following the instructions generated by the SMRT Link software (PacBio, version 10) and the ‘HiFi Reads’ application. The HiFi library was sequenced on 8M SMRT™ Cells for 30 hours with the Sequel^®^ II Binding 2.2 and the Sequel^®^ II Sequencing 2.0 chemistry on the Sequel^®^ II.

##### 10x genomic linked read sequencing

Ultra-long gDNA from species *Apus apus*, *Ptilotula penicillata*, and *Loriculus galgulus* was used for 10x genomic linked read sequencing following the manufacturer’s instructions (10X Genomics Chromium Reagent Kit v2, revision B). Briefly, 1 ng of ultralong gDNA was loaded into 10X Genomics GEM droplets (Gel Bead-In-EMulsions) making use of the Chromium device. gDNA molecules were amplified in these individual GEMS in an isothermal incubation using primers that contain a specific 16 bp 10x barcode and the Illumina R1 sequence. After breaking the emulsions, pooled amplified barcoded fragments were purified, enriched and used forIllumina sequencing library preparation, as described in the protocol. Sequencing was done on a NovaSeq 6000 S1 flow cell using the 2x150 cycles paired-end regime plus 8 cycles of i7 index.

##### Bionano optical mapping

For *Apus apus*, *Ptilotula penicillata*, and *Loriculus galgulus*, optical mapping was done with the Bionano Prep Direct Label and Stain DLS DNA Kit (catalog #8005, Bionano Genomics, San Diego), according to the manufacturer’s Protocol (Document Number 30206, Document Revision F). Briefly, 750 ng of ultra-long gDNA was fluorescently labeled at defined sequences, making use of the nicking-free Bionano Direct Label Enzyme (DLE-1). For further visualization, the DLE-1 labeled gDNA backbone was stained with DL-Green. Labeled molecules were imaged using the Bionano Saphyr system. Data were generated from one Bionano flow cell with a total yield of 341, 997, 215 and 320 Gb, respectively, in molecules larger than 150 kb.

##### Chromatin conformation capturing (HiC)

Chromatin conformation capturing was done using the Arima-HiC kit (Article Nr. A410109) for *Ptilotula penicillata* and ARIMA-HiC High Coverage Kit (Article Nr. A101030-ARI) for *Apus apus*, and *Loriculus galgulus*, following the Arima documents (User Guide for Animal Tissues, Part Number A160132 V01 and Part Number A160162 v00; Guide to Preparing Crosslinked Nucleated Blood). Briefly, 66 mg flash-frozen powdered muscle tissue from *Ptilotula penicillata*, 46 mg muscle tissue in ethanol from *Nymphicus hollandicus*, 50 µl pure blood sample from *Apus apus* and 50 µl blood preserved in ethanol from *Loriculus galgulus* were crosslinked chemically. All the tissue and blood samples that were stored in ethanol were first treated with ethanol removal steps (3 times washing in 1 ml of 1 x PBS buffer) before the crosslinking procedure. The crosslinked genomic DNA was digested with a cocktail of two (*Ptilotula penicillata*) or four (*Apus apus* and *Loriculus galgulus*) restriction enzymes. The 5’-overhangs were filled in and labeled with biotin. Spatially proximal digested DNA ends were ligated, and the ligated biotin containing fragments were enriched and used for Illumina library preparation, following the ARIMA user guide for Library preparation using the Kapa Hyper Prep kit (ARIMA Document Part Number A160139 v00). The barcoded HiC libraries were run on an S4 flow cell of an Illumina NovaSeq 6000 with 2x150 cycles.

### Genome assembly

#### Apus apus, Ptilotula penicillata, Nymphicus hollandicus, and Loriculus galgulus

The genomes of *Apus apus*, *Ptilotula penicillata*, and *Loriculus galgulus* were assembled using PacBio Continuous Long Reads (CLR), 10X Genomics linked-reads, Bionano DLS optical maps and Arima Chromatin Conformation Capture (HiC) libraries as follows. Initial contigs were assembled from PacBio subreads using Falcon (v1.8.1) and Falcon Unzip (v1.3.7) (*121*). Remaining haplotypic duplications in the primary contig set were then removed using purge-dups (v.0.0.3) (*122*). The genome of Nymphicus hollandicus was assembled using PacBio HiFidelity (HiFi) Circular Consensus Sequences (CCS) and Arima HiC libraries.

Scaffolding was performed using 10x genomics linked Illumina reads, Bionano optical maps, and HiC chromosome conformation capture Illumina read pairs. To this end, 10x reads were first mapped to the contigs using Long Ranger (v2.2.2) and scaffolded using Scaff10X (v4.2). The 10x linked-reads were then re-mapped to the scaffolded assembly produced by Scaff10X and any false joins were removed using the Break10X command from Scaff10X. Next, we used optical maps from Bionano DLE1-labeled DNA molecules. The Bionano assembly was produced from optical-mapped reads using Bionano Solve Assembly (v3.5.1) with parameters “non-haplotype” and “no-Extend-and-Split”. The initial scaffolds from Scaff10X were then further scaffolded using Bionano Solve Hybrid Scaffold (v3.5.1). HiC reads were then mapped to the resulting scaffolds using bwa-mem (v0.7.17-r1188) (*123*, *124*). We then followed the VGP scaffolding pipeline (https://github.com/VGP/vgp-assembly/tree/master/pipeline/salsa) (*31*), which uses salsa2 (v2.2) (*125*). Finally, we manually curated the scaffolds to join those contigs missed by salsa2 and break those joins which were spuriously created. In order to visually evaluate the scaffolding, reads were again mapped to the scaffolded assembly using bwa-mem, filtered and deduplicated using pairtools (v0.3.0) and the multi-cooler file for higlass (v0.4.17) was produced using cooler (v0.8.11). We then curated the assembly through an iterative process of joining together scaffolds which were missed by the automated scaffolding software and breaking any false joins between two chromosomes until no further corrections could be made to the assembly.

After scaffolding, we closed assembly gaps using the PacBio CLR read data. To this end, we mapped the original subreads.bam files to the scaffolded assembly using pbmm2 (version 1.3.0) with arguments “--preset SUBREAD -N 1”. For the assembly of Apus apus, we then used Dentist (v1.0.0) (*126*). For the assemblies of *Ptilotula penicillata* and *Loriculus galgulus* we used gccp (v2.0.2) to close gaps. Based on the read-piles created by reads spanning across gap regions, we can create a consensus sequence to replace the N sequences in our genome. We used gcpp to polish the gap regions and their 2 kb upstream and downstream flanks. We then replaced the assembly gap and its flanking region with those regions that were polished with high-confidence consensus sequence (no N’s remaining in the polished sequence and no lower-case a/c/g/t).

To increase base level accuracy, we polished the resulting assembly using the 10X linked-reads as done in the VGP pipeline (https://github.com/VGP/vgp-assembly/tree/master/pipeline/freebayes-polish) (*31*). Briefly, 10X reads were mapped to the genome using Long Ranger (v2.2.2) and variants were called using freebayes (v1.3.2) with argument “-g 600” to ignore regions with coverage over 600X. The detected variants were filtered using bcftools (v1.12-21-ga865a16) for variants with quality score greater than 1 and genotype of homozygous alt (AA) or heterozygous (Aa) using the command bcftools view -i ‘QUAL>1 && (GT=“AA” || GT=“Aa”)’. The consensus was then called using the command bcftools consensus -i’QUAL>1 && (GT=“AA” || GT=“Aa”)’ -Hla, which takes the longest allele in heterozygous cases. This procedure was performed twice. Finally, we used Merqury (v1.0) (*127*) with the 10x data. For the assembly of *Loriculus galgulus* we performed further polishing into two haplogroups. Chromosomes were phased by applying an adapted version of the DipAsm pipeline (*128*), calling heterozygous sites using 10x reads and creating phase blocks by linking heterozygous sites in 10x, Hi-C and PacBio reads using Hapcut2 (git commit 1ee1d58) and whatshap (v1.6). The whatshap-tagged PacBio and 10x reads, binned into H0 and H1, and H0 and H2, were then used to haplotype-polish the chromosomes. Arrow was run using the combined H0 and H1 haplotagged PacBio CLR reads, followed by freebayes polishing using the combined H0 and H1 haplotagged 10x reads as above. Correspondingly the H2 and H0 tagged reads were used for the second haplotype.

#### Florisuga fusca

The genome of *Florisuga fusca* was assembled from Illumina short reads (see above) using a hybrid approach similar to (*129*). Initial assembly was conducted with IDBA-UD (*130*), followed by further assembly using CELERA ASSEMBLER v8.1 (*131*), supported by subsets of the short-read data. Scaffolds were ordered using SSPACE2 (*132*) and aligned to the galGal4 assembly via RAGOUT v1 (*133*). Gap filling was performed with GAPCLOSER (*134*) to enhance contig N50.

#### Acanthiza pusilla and Phylidonyris novaehollandiae

We assessed the quality of the sequencing data generated by the Illumina HiSeq 2500 platform using FastQC (*135*). Adapters were removed using Trimmomatic (*136*). Genome assembly was performed with AllPaths-LG v52488 (*137*).

#### Furnarius rufus and Trichoglossus moluccanus

De novo genome assembly for both species was performed using the nf-core/neutronstar analysis pipeline implemented in Nextflow (*138*). Supernova v2.1.0 (*139*) was used to construct initial assemblies from Illumina NovaSeq 6000 sequencing data derived from 10X Genomics libraries. For *Trichoglossus moluccanus*, additional scaffolding was performed using the 3D-DNA pipeline (*140*), leveraging Hi-C data to improve assembly contiguity.

### Annotation

#### Modeling and masking repeats

For the newly generated assemblies, we used RepeatModeler (*141*) to identify repeat families and RepeatMasker (*142*) to soft-mask repeats. For other bird genomes, we used the available repeat masker annotations provided by NCBI. All species names and their assemblies are listed in table S1.

#### Generating pairwise genome alignments

To compute pairwise genome alignments between chicken (galGal6 assembly), *T*. *guttata* (GCF_003957565.2) and *P*. *major* (GCF_001522545.3) as references and other birds as query species, we used LASTZ version 1.04.00 (*143*) with parameters K = 2,400, L = 3,000, Y = 9,400, H = 2,000 and the default scoring matrix, axtChain (*144*), chainCleaner (*145*), and RepeatFiller (*146*) (all with default parameters). For the downstream analyses except TOGA, we excluded chains with a score less than 100,000.

#### Protein-coding gene annotation

To achieve a high completeness of the protein-coding gene annotations, we combined four different types of evidence: 1) homology-based predictions, 2) protein data, 3) transcriptomic data, and 4) ab initio gene predictions.

First, we generated pairwise alignments between all target genomes and three reference genomes: *G. gallus* (chicken), *T. guttata* (zebra finch), and *P. major* (great tit). We then used TOGA 1.0 (*34*) with default parameters to project transcripts annotated for reference species onto the target genomes. As reference annotations, we used NCBI annotations of *T. guttata* (46,022 transcripts) and *P. major* (41,530 transcripts). For *G. gallus* we combined NCBI annotation with APPRIS principal isoforms (resulting in 64,081 transcripts).

Second, we prepared protein evidence aligning translated protein data to the target genomes using GenomeThreader (v1.7.0) (*147*) applying the Bayesian Splice Site Model (BSSM) trained for chicken and default parameters aside from those specified below. For protein alignments, we used parameters: a seed and minimum match length of 20 amino acids (preseedlength 20, prminmatchlen 20) and allowed a Hamming distance of 2 (prhdist 2). For the transcript alignments, we used parameters: a seed length and minimum match length of 32 nucleotides (seedlength 32, minmatchlen 32). At least 70% of the protein or mRNA sequence was required to be covered by the alignment (-gcmincoverage 70), and potential paralogous genes were also computed (-paralogs).

Third, for five species (black jacobin, New Holland honeyeater, rainbow lorikeet, brown thornbill, and rufous hornero), we used available transcriptomic data obtained from (*148*) as an additional source of evidence and assembled transcripts with Stringtie (v2.1.2).

Fourth, we ran de novo gene prediction using Augustus (v3.3.3) (*149*) providing TOGA projections, mapped protein and RNA-seq data as hints. The bam2hints module of Augustus with the introns-only flag was used to produce RNA-seq hints. Priority 4 was given for RNA-seq hints. With the script align2hints.pl (provided with BRAKER) protein hints from GenomeThreader alignments were prepared. Priority of 4 was given to protein-derived hints. Finally, RNA-seq- and protein-derived hints were merged using the join_mult_hints.pl script from Augustus. Prediction of additional splice sites was enabled with the allow_hinted_splice_sites flag (--allow_hinted_splicesites=gcag,atac). Prediction of untranslated regions was disabled (--UTR=off). The resulting files were merged using the Augustus joingenes module. bedtools intersect was used to exclude all Augustus predictions having more than 10% overlap with a repeat region. Next, all predictions were converted to proteins and queried with blastp against the Swissprot database with an E-value cut off 1e-10. An in-house script was used to filter blast results allowing only hits matching a sequence in the vertebrates database or having lengths greater than 200 amino acids (<u>filter_fasta_with_blast.py</u>).

After generating the four types of gene evidence, we integrated them into a single annotation per species. To this end, we generated a set of consensus transcripts using EVidenceModeler (v1.1.1) (*150*), providing combined aligned protein and RNA-seq data, TOGA projections, and de novo transcripts as input. We assigned weights prioritizing TOGA projections over protein data and RNA-seq data followed by de novo predictions (8, 2, 2, and 1 respectively). Since EVidenceModeler predicts only one consensus gene model per locus, the final set of gene models lacks information about 1) nested genes located in the intronic regions of other genes; 2) alternative isoforms. To rescue those, we added high-confidence gene predictions to the resulting set of consensus models by adding TOGA transcript projections that were identical for at least two out of the three reference species, as done before (*32*). The final annotation comprised between 33,507 (sun parakeet) and 131,520 (white-plumed honeyeater) transcripts and between 19,321 (sun parakeet) and 44,121 (rainbow lorikeet) genes (table S1).

Since these annotations only contain coding exons, we subsequently added UTR exons using available RNA-seq data. We used a custom script (<u>cds_add_utrs_from_stringtie.py</u>) to extend gene models into UTR regions based on the RNA-seq Stringtie assembly. The donor site of the first intron or the acceptor site of the last intron of a predicted transcript was required to match perfectly by RNA-seq structure. For single-exon transcripts, the exon start and end were required to lie within the corresponding sequence evidence. If there were conflicts between several Stringtie models, the UTR regions were assigned based on the majority.

For the species with the available NCBI annotation, we downloaded the available annotations. We assessed the completeness of all annotations (NCBI and ours) using the BUSCO pipeline (v5.2.1) (*28*) with the set of 8338 conserved, single-copy avian genes.

### Positive selection analysis

To identify the one-to-one orthologs, we used TOGA (*34*) based on the chicken gene annotation, keeping only projections classified as intact, partially intact or uncertain loss. Next, we aligned identified orthologs with MACSE_v2 (*151*) with default parameters, exon by exon, taking into account the codon structure of a gene. We filtered the alignments with HmmCleaner (v0.180750) (*152*) and then applied additional filtering with an in-house script (<u>manual_filter_msa.py</u>), removing sequences entirely if less than 50% of the sequence was aligned and removing codon alignment columns if less than 50% of species were aligned. We inspected final alignments manually for potential misalignments. We analyzed positive selection with the aBSREL package of HyPhy (v2.5.58) (*35*, *153*): for each transcript we ran four models: 1) the default model; 2) a model enabling synonymous rate variation (--srv Yes); 3) a model enabling correction for multinucleotide substitution (--multiple-hits Double+Triple); and 4) a model enabling both SRV and multihits (--multiple-hits Double+Triple --srv Yes). For every model we enabled tolerating numerical errors (ENV=’TOLERATE_NUMERICAL_ERRORS=1). For each transcript, we calculated the weighted p-value based on AICc (Akaike information criterion corrected) values using the formula from (*99*).

For each gene to be under positive selection at a specific branch, we required that at least ⅓ of transcripts annotated for this gene to have signal of positive selection. For each gene we assigned a ‘relevance ratio’ based on the ratio of the number of sugar branches to the total number of branches with a significant signal of positive selection (p-value < 0.01 corrected for the branch number); branches classified as ambiguous were not included in either count. Genes having the relevance ratio = 1.0 were shortlisted and additionally examined for pathway enrichment.

We performed the enrichment analysis with the R package ClusterProfiler (v4.8.2) (*154*) using the human OrgDb org.Hs.eg.db and Gene Ontology (GO) Biological Processes subontology. To be able to use the human database, we converted chicken gene IDs into human gene IDs with Biomart (https://useast.ensembl.org/biomart/martview). The entire genome was considered as the background gene set. We selected significantly enriched terms applying FDR cutoff=0.3.

To test gene-level repeatability, we used a two-sided Fisher’s exact test. We compared the ratio of the number of genes showing repeated (i.e. shared between at least two groups) vs. lineage-specific positive selection in sugar-feeders with the ratio calculated for a core non-sugar control group.

To identify specific sites under positive selection in selected gene candidates - specifically, in *MLXIPL* and *HK3* genes - we used the branch-site model of codeml from the PAML (v4.9) package (*155*), selecting the ancestral honeyeater branch (for *HK3*) and all sugar-consuming branches (for *MLXIPL*) as the foreground. Sites under positive selection were defined as sites having Bayes Empirical Bayes (BEB) posterior probabilities >0.5.

### Homology modeling and protein structure manipulation

We performed homology modeling using the swissmodel.expasy.org modeling service (*156*) providing the New Holland honeyeater sequence. Since no crystal structure of hexokinase 3 was solved, we used a human hexokinase 2 crystal structure (PDB ID: 5HEX) as a template. The structure visualization and manipulation were done in ChimeraX-1.9 (*157*).

### Convergent amino acid substitutions in protein-coding sequences

We utilized CSUBST v1.0.0 (*36*) to identify convergent amino acid substitutions in protein-coding sequences. CSUBST employs the ω_c_ metric, a ratio of non-synonymous to synonymous combinatorial substitution rates. This metric is designed to be robust against phylogenetic errors and to discern genuine adaptive convergence from false positives. Unlike the traditional ω measure, ω_c_ contrasts the ratio of observed to expected non-synonymous combinatorial substitutions with the synonymous counterpart between distinct phylogenetic branches. For convergence in a pairwise branch comparison, we set criteria: ω_c_ ≥ 3, O_c_^N^ ≥ 3 (any2spe category).

### CNEE alignments

We obtained vertebrate conserved non-exonic elements (CNEEs) from previously published sources (*37–40*). These sources included: 578,464 CNEEs from Lowe et al. (*38*), identified from 5 vertebrates (human, mouse, cow, medaka, and stickleback); 1,044,557 CNEEs from UCSC (*40*), identified from 5 vertebrates (human, mouse, rat, chicken, and Fugu rubripes); 1,954,198 CNEEs from Sackton et al. (*39*), identified from 42 species (35 bird species, including 10 paleognath and 23 neognath genomes, plus 7 non-avian reptilian outgroups); and 1,111,278 CNEEs from Craig et al. (*37*), identified from 23 sauropsid species. These were then lifted over to *Gallus gallus* (galGal6 or GRCg6a assembly) coordinates using halLiftover (*158*).

To ensure the CNEEs met the general expectations for length and conservation, we performed a series of cleaning steps. Initially, we eliminated elements exceeding 5 kb in length and merged elements that overlapped or were within 5 bp of each other. Elements shorter than or equal to 50 bp were also removed. Subsequently, we excluded elements overlapping with any exonic regions in GRCg6a. Lastly, we discarded CNEEs that were not conserved as defined by phyloP (*159*) using the CON mode and LRT method, with adjusted p-values greater than 0.1, ensuring a baseline level of conservation across the species included in our whole-genome alignment.

Following the pipeline established in (*39*), we generated CNEE alignments from the curated set of CNEEs. In brief, we extracted sequences for the curated CNEEs from all species included in the whole-genome alignment. We used the halLiftover tool from HAL tools v2.1 to generate psl files, which mapped chicken reference coordinates to each target species. Custom Perl scripts (*39*) were then used to convert halLiftover output into FASTA format files. Sequences for individual CNEEs were omitted if the target region was duplicated in a given taxon or exceeded twice the reference chicken length.

Individual CNEEs were aligned using MAFFT v7.304 (*160*) with the ‘ginsi’ option. The alignments were then trimmed using TrimAl v1.4.rev22 build[2015-05-21] and the ‘-noallgaps’ option, which removes columns composed entirely of gaps. Subsequently, the trimmed fasta files were merged into a single file using catsequences (*161*), with the provision of a partition file defining the start and end positions of each locus. We obtained 363,476 CNEE alignments. All species had a liftover rate exceeding 96% relative to the curated set of CNEEs in the GRCg6a coordinates. Furthermore, the majority (74.6%) of the CNEE alignments contained sequences from all species. While we did not exclude CNEEs based on species representation at the alignment stage, in the subsequent PhyloAcc analysis, we required elements to be present in at least 80% of the conserved species (31 non-target species), resulting in 363,213 CNEEs being analyzed for accelerated evolution.

To further validate the conservation status of our CNEEs and address potential biases in element selection, we conducted a permutation analysis (500 iterations) using the phyloP score based on ancestral repeats associated with the 363-bird genome paper (*30*). We calculated the average of these scores across each CNEE and compared them to randomly shuffled control regions with matching length distributions (avoiding exonic overlaps). The empirical p-value of 0.002 confirms that our CNEEs are significantly more conserved than expected by chance, supporting that our selection process is robust and not biased by the composition of reference species. This validation provides additional evidence that the CNEEs we analyzed represent genuinely conserved elements across the avian phylogeny.

### PhyloAcc

We employed PhyloAcc v2.2.0 (*41*, *162*), a Bayesian method designed to model shifts in substitution rates within conserved elements across a phylogeny. Our objective was to determine whether a CNEE displays accelerated substitution rates in comparison to a neutral model across each branch of the avian phylogeny. Although accelerated CNEEs could result from either positive selection for new regulatory function or relaxed constraint on ancestral elements, both mechanisms indicate regulatory divergence likely relevant to the physiological adaptations required for sugar-feeding.

To derive a neutral model, we first extracted fourfold degenerate sites from the entire genome alignment using the PHAST software package (*163*). Subsequently, we utilized the ‘phyloFit’ program (*164*) to estimate a neutral model from these sites. We performed a PhyloAcc analysis specifically for the sugar-consuming birds, designating these species as ‘TARGETSPECIES’, using the species tree run mode. Further program settings are detailed on our GitHub repository (https://github.com/maggieMCKO/nectargenomics/). To identify sugar-specific acceleration, we conducted an additional PhyloAcc run (referred to as the control run), setting the core non-sugar-consuming birds (defined above), as ‘TARGETSPECIES’.

### Defining accelerated CNEEs

We used PhyloAcc (*41*, *162*) to identify genomic elements exhibiting lineage-specific acceleration. This method relies on sequence alignments across a phylogeny to model substitution rate shifts using Bayes factors. PhyloAcc employs three nested models: 1) the Null model (M0), which assumes no shift in substitution rates; 2) the lineage-specific model (M1), which permits rate shifts in predefined lineages, such as sugar-consuming birds.; and 3) the full model (M2), which allows rate shifts in any branch. To detect CNEEs with lineage-specific accelerations, we compared the marginal likelihoods of these models, defining three Bayes factors: Bayes factor 1 (BF1) = (P(Y|M_1_))/(P(Y|M_0_)) and Bayes factor 2 (BF2) = (P(Y|M_1_))/(P(Y|M_2_)).

We identified accelerated CNEEs using PhyloAcc with two sets of criteria with increasing specificity to sugar-consuming lineage acceleration. For the loose set, we applied a cutoff of logBF1 > 10 and a ‘relevance ratio’ > 0.5, which is conceptually similar to the ratio used for positively selected genes. For each CNEE, the relevance ratio was calculated as the number of sugar branches, including internal branches, in the accelerated state (posterior probability under the full model > 0.9) divided by the number of background branches (excluding ambiguous branches) in the accelerated state. This approach identifies CNEEs with evidence of acceleration predominantly in sugar-consuming lineages relative to background evolution.

For the strict set, we applied more stringent criteria of logBF1 > 10 and logBF2 > 3. The logBF2 > 3 threshold requires acceleration in sugar-consuming or core non-sugar consuming lineages to be substantially greater than in other lineages across the phylogeny, providing more stringent evidence for lineage-specific regulatory evolution associated with dietary specialization.

To categorize accelerated CNEEs within specific sugar-consuming or core non-sugar groups, we required at least one branch in each group to have a posterior probability exceeding 0.9 for being in the accelerated state.

Comparison of GO enrichment analysis results (see below) between loose and strict sets revealed that although the strict set identified substantially fewer accelerated CNEEs and consequently fewer significantly enriched GO terms, the fold enrichment values increased markedly in the strict set (fig. S6A and B). Therefore, we used the strict set for all downstream analyses including enrichment tests, Fisher’s exact tests for repeated evolutionary changes, and T2D enrichment analyses.

### CNEE statistical analysis

We assessed whether repeated evolutionary changes in regulatory elements are more frequent in sugar-consuming lineages using Fisher’s exact tests. For our main statistical test using the strict CNEE set, we performed a two-sided Fisher’s exact test with a 2×2 contingency table design. CNEEs were categorized as either accelerated or not accelerated, and lineages were classified as sugar-consuming or core non-sugar controls. This approach directly tests whether the proportion of accelerated CNEEs differs significantly between dietary groups.

To provide detailed insights into patterns of repeated changes across multiple lineages (reported in supplementary figures), we conducted additional analyses examining pairwise, triple, and quadruple combinations. Additionally, for the gene-level repeatability analysis, we utilized GREAT (see below) to associate CNEEs with genes and performed a two-sided Fisher’s exact test.

We employed the GREAT software (*42*) to associate CNEEs with genes, using its ‘createRegulatoryDomains’ utility set to the default ‘basalPlusExtension’ configuration. This configuration establishes a ‘regulatory domain’ for each protein-coding gene, extending 5 kb upstream and 1 kb downstream from the transcription start site. Furthermore, the regulatory domain of each gene is expanded to the basal regulatory regions of the nearest upstream and downstream genes, with a maximum extension limit of 1 Mb in each direction. This allows for the incorporation of distal binding sites into its analysis, thereby providing a broader perspective on the potential regulatory influences of CNEEs on gene expression.To enhance the annotation of chicken genes (GRCg6a) for the Gene Ontology (GO) and Kyoto Encyclopedia of Genes and Genomes (KEGG) by leveraging human databases, we created custom GO and KEGG databases. We used the Biomart library (v.2.56.0, Ensembl ver106) in R to map chicken Entrez gene IDs to their corresponding Ensembl and human Ensembl gene IDs, based on Ensembl’s ortholog database.

For GO annotation, we employed the Biomart library to retrieve GO IDs for each human gene, thereby increasing the number of chicken genes linked to GO IDs from 2,203 to 7,586. For KEGG pathways, we converted human Ensembl gene IDs to Entrez IDs using the ‘org.Hs.eg.db’ R library (v.3.17.0) and then used the ‘KEGGREST’ R library (v.1.40.0) to identify associated KEGG pathways. This expanded the number of chicken genes linked to KEGG pathways from 3,482 to 6,680.

To focus on biologically meaningful enrichments and reduce computational noise, we first filtered our analysis to include only GO terms with at least 40 genes associated with CNEEs. For these filtered GO terms, we conducted hypergeometric tests to assess enrichment while accounting for the non-random genomic distribution of CNEEs, which tend to cluster in specific loci. We also calculated fold enrichment for each GO term using the formula: fold enrichment = (k/n)/(K/N), where k is the number of accelerated CNEEs associated with the GO term, n is the total number of accelerated CNEEs, K is the total number of CNEEs associated with the GO term, and N is the total number of CNEEs. In our final results, we only show significantly (FDR-adjusted p-value < 0.3) enriched GO terms that had at least 3 genes from input gene lists. Additionally, we exclude enriched pathways that are shared by at least 2 control groups. This combined approach of pre-filtering, hypergeometric testing, and post-test filtering ensures a rigorous identification of functionally relevant enrichments while accounting for the non-random distribution of CNEEs across the genome.

### Tissue collection and RNA sequencing

Tissues were excised, each piece was cut into aliquots of approximately 100 mg and then transferred into a Dewar with liquid nitrogen for several minutes, or placed on dry ice, or placed in 5-10 volumes of RNAlater (Thermofisher Scientific, AM7020) solution and incubated overnight at 4°C before storage at -80°C. Throughout the whole process, an RNAse-free environment was ensured and all surfaces and instruments were treated with RNase Zap solution (Thermofisher Scientific, AM9780).

Total RNA was isolated using the RNeasy mini kit (Qiagen). The concentration and the RNA integrity number (RIN) were analyzed using the Bioanalyzer (Agilent). RNA-seq library preparation and the sequencing analysis were performed in the Dresden-Concept Genome Center (DcGC; TU Dresden, Germany). In detail, mRNA was enriched from 400 ng total RNA by poly-dT enrichment using the NEBNext Poly(A) mRNA Magnetic Isolation Module according to the manufacturer’s instructions. The polyadenylated RNA fraction was eluted in 11.5ul 2x first-strand cDNA synthesis buffer (NEBnext, NEB). After chemical fragmentation, by incubating for 13 min at 94°C the samples were directly subjected to the workflow for strand-specific RNA-seq library preparation (Ultra II Directional RNA Library Prep, NEB). In short, the first and second strand synthesis followed an end repair and A-tailing of the fragments that were ligated with the universal NEB hairpin loop adapter. After ligation, adapters were depleted by an XP bead purification (Beckman Coulter) adding the beads solution in a ratio of 1:0.9. During the PCR enrichment with 12 cycles, unique dual index primers were incorporated carrying the sequence for i7 and i5 tails (Primer 1: AAT GAT ACG GCG ACC ACC GAG ATC TAC AC XXXXXXXX ACA TCT TTC CCT ACA CGA CGC TCT TCC GAT CT, Primer 2: CAA GCA GAA GAC GGC ATA CGA GAT XXXXXXXX GTG ACT GGA GTT CAG ACG TGT GCT CTT CCG ATC T; X represents the different barcode sequences). After two more AMPure XP bead purifications (in a ratio of 1:0.9), libraries were quantified using the Fragment Analyzer (Agilent) and sequenced on an Illumina NovaSeq 6000 with 2x 100 bp reads using an S4 flowcell to an average depth of 30 mio. read pairs.

## Analysis of transcriptomic data

We trimmed adapters and low-quality sequences from the reads using Trimmomatic (v0.39) with parameters: PE ILLUMINACLIP:****:2:30:10:2:keepBothReads LEADING:3 TRAILING:3 SLIDINGWINDOW:4:15 MINLEN:36. We quantified transcript-level expression with Kallisto (v 0.46.1) with default parameters and bootstrap 100 (-b 100). To run Kallisto, we generated species-specific transcriptome indexes based on our generated annotations. We quantified gene-level expression with tximport R package. We ran differential gene expression analysis with DESeq2 (R v4.1.1), specifying contrasts to perform between species pairwise and multifactor cross-tissue analyses. Each test is limited to a single phylogenetically independent pairwise comparison between the sugar-consuming target species and its matched non-sugar-consuming control. We performed the enrichment analysis with the R package ClusterProfiler (v4.8.2) (*154*) using human OrgDb ‘org.Hs.eg.db’ and Gene Ontology (GO) Biological Processes subontology. To be able to use the human database, we converted chicken gene IDs into human gene IDs with Biomart (https://useast.ensembl.org/biomart/martview). For the pairwise comparisons, we specified the background set to only list genes having a non-zero expression in at least one of the samples in the tested tissue. For the cross-tissue comparison, we specified the background set as genes having a non-zero expression in either liver or pectoralis. We selected significantly enriched terms applying FDR cutoff=0.3. Finally, we removed redundancy of the raw GO term lists excluding direct children terms - more specific instances of broader parent terms in the GO hierarchy - of the terms that were already present in the list.

### Manual curation of hexokinase 3 and *MLXIPL* gene models

Reference exons from hexokinase 3 (*HK3*), *MLXIPL*, and *MLX* gene predictions were downloaded from NCBI (for *HK3*: chicken (XM_001231328.5), zebra finch (XM_030283802.2), great tit (XM_015642097.2), lance-tailed manakin (XM_032703318.1), and human (NM_002115.3); for *MLXIPL*, chicken (NM_001398117.1); for *MLX* chicken (NM_001030930.1) and Anna’s hummingbird (XM_030465864.1)) and mapped to genome assemblies (for HK3, to our New Holland honeyeater assembly, as well as to the publically-available superb lyrebird assembly (GCA_013396355.1) and rufous treecreeper (GCA_013398175.1); *MLXIPL* and *MLX* were curated from our common swift assembly, as well as from the publically-available Anna’s hummingbird assembly (GCA_003957555.2)). Transcriptomes were assembled using HISAT2 v2.1.0 and STRINGTIE v.2.2.1 (New Holland honeyeater: liver, duodenum, heart, palate, and pectoralis samples; common swift: duodenum samples; and Anna’s hummingbird: duodenum and caudal intestine samples) to validate the predicted transcripts.

There are several *MLXIPL* isoforms annotated for Anna’s hummingbird (fig. S12C), so we manually curated the most likely functional isoforms based on the transcriptomic data. We found that two isoforms have a dominant expression (fig. S12D). These isoforms differ only by 279 base pairs in the first exon, the long isoform has an upstream start codon, while the short isoform uses an alternative downstream start codon (fig. S12C,D). Both variants were expressed in the duodenum, with the shorter variant also observed in one of the caudal intestine samples. The domain structure shows that the short isoform potentially lacks a part of a functional (LID) domain. Additionally, the short isoform possibly lacks a bird-specific nuclear localization signal (NLS) (fig. S12E). Similar, corresponding variants were observed in the common swift genome. The identified gene versions for *MLXIPL*, *MLX* from swift and hummingbird, and *HK3* from honeyeater, lyrebird, and human were synthesized by Genewiz for further experimental validation.

### Hexokinase purification

The synthesized *HK3* genes from New Holland honeyeater, rufous treecreeper, superb lyrebird, and human were cloned into the pCoofy6 vector (*165*) using the SLIC cloning method (*166*). This created fusion proteins with His6-Sumo3 tags at the N-terminus of HK3, facilitating purification. The *E*. *coli* expression system was chosen based on previous successful purifications of human hexokinases (HK2 and HK3) (*167*, *168*).

Recombinant HK3 proteins were expressed in *Escherichia coli* Rosetta (DE3) T1 (Novagen N59205). Cultures were grown in auto-induction medium (*169*) at 37°C for 2 hours, in the presence of appropriate antibiotics. The temperature was then lowered to 20°C and cells were grown overnight. Cells were harvested by centrifugation at 6,000 rpm at 4°C for 10 minutes. The cell pellet was resuspended in lysis buffer (50 mM Sodium Phosphate, pH 8.0, 500 mM NaCl, 10 mM imidazole, 10% Glycerol, 1 mM TCEP, 0.1% protease inhibitor cocktail, and 2.4 U/ml Benzonase). Cells were lysed by LM20 microfluidizer, and the lysate was clarified by centrifugation at 20,500 rpm for 30 minutes at 4°C. The supernatant was loaded onto a Nickel-Chelating Resin pre-equilibrated with binding buffer (50 mM Sodium Phosphate, pH 8.0, 500 mM NaCl, 10 mM imidazole, 10% Glycerol and 1 mM TCEP) at 4°C. The resin was washed with binding buffer followed by washing buffer (same as binding buffer but with 20 mM imidazole). The His-tagged HK3 was eluted with elution buffer (50 mM Sodium Phosphate, pH 8.0, 500 mM NaCl, 250 mM imidazole, 10% Glycerol and 1 mM TCEP). The eluted His-SUMO-HK3 was digested with SenP2 at 4°C in dialysis buffer (same as binding buffer) overnight. The Ni-NTA column was used again to remove His-SUMO, undigested His-SUMO-HK3, and SenP2 protease (in which the hexahistidine tag was also fused to the N-terminal), thus resulting in the native HK3. HK3s were further purified by size-exclusion chromatography using a HiLoad Superdex 200 column equilibrated with filtration buffer (50 mM Tris pH 7.4, 250 mM NaCl, 5% Glycerol and 1 mM TCEP). The purified protein concentration was determined by A280 measurement. Protein purity was assessed using SDS-PAGE. Finally, the purified HK3 in storage buffer (50 mM Tris, pH 7.5, 250 mM NaCl, 5% Glycerol, 1 mM TCEP) was snap-frozen in liquid nitrogen and stored at -80°C until further use.

### Hexokinase functional assay

The purified HK3 enzymes from New Holland honeyeater, rufous treecreeper, superb lyrebird, and human were diluted to a final concentration of 200 nM for activity measurements. The molecular weights of HK3 were 112,470 Da (New Holland honeyeater), 112,620 Da (rufous treecreeper), 112,850 Da (superb lyrebird), and 109,905 Da (human), corresponding to mass concentrations of 22.494 µg/mL, 22.524 µg/mL, 22.57 µg/mL, and 21.981 µg/mL, respectively. To determine kinetics, hexokinase activity was assessed using the Abcam hexokinase activity assay kit (ab136957) according to the manufacturer’s instructions, except the HK substrate was replaced by varying glucose concentrations (1, 0.5, 0.15, 0.075, 0.0375, 0.01875, and 0.009375 mM). The assay was performed at 25°C using a Flexstation3 microplate reader (Molecular Devices), with optical density (OD) readings taken every 30 seconds at 450 nm. Sample sizes were between 7 and 13 for each enzyme and glucose concentration combination. Kinetic parameters were estimated using R statistical software. Haldane’s formula for substrate inhibition (*170*) was applied to fit the data:

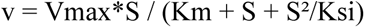

where v is the reaction velocity, S is the substrate concentration, Vmax is the maximum velocity, Km is the Michaelis constant, and Ksi is the substrate inhibition constant. The nonlinear least squares function in R was used to fit the model to the experimental data, with the PORT algorithm for optimization. Vmax, Km, and Ksi values were extracted from the model fit, and 95% confidence intervals were calculated, characterizing the hexokinase activity for HK3 enzymes (fig. S10). Statistical significance between species for each parameter was determined by non-overlapping 95% confidence intervals.

### Histological analysis

Tissues were excised, each piece was cut into aliquots of approximately 100 mg. Tissue aliquots were fixed in 4% PFA solution at 4°C overnight. The next day, tissue aliquots were placed in fresh sterile PBS and shipped with cooling packs at 4°C. The fixed tissues were stored in PBS for 48 hours. Next, tissues were incubated in 10% sucrose solution (24 hours) followed by incubation in 30% sucrose solution (24 hours). Then tissues were covered with OCT (Optimum Cutting Temperature compound, Thermofisher Scientific, 23730571) and placed in a labeled cryomold. Isopentane (2-methylbutane) was chilled in a steel beaker for 3-5 minutes using liquid nitrogen until white precipitates started to form on the bottom and walls of the beaker. The cryomolds with tissues were immersed in chilled isopentane for 6-12 seconds (depending on the tissue size). Cryomolds with tissues were transferred on dry ice for 15 mins to let isopentane evaporate. Cryomolds were filled up to the top with OCT and stored at -80°C until sectioning. Sectioning was performed on the cryostat NX70 (Thermofisher Scientific) with the following settings: -15°C temperature for the tissue and -29°C for the knife, 10m sections in a series of 2. The sections were collected on superfrost slides (Langenbrinck) and stored at -20°C until staining. Cryosections were stained for lipids with Oil Red O histological staining. In summary, sections were dried for 1 hour at room temperature. Next, slides were incubated in 60% isopropanol for 20 sec followed by the incubation in Oil Red O solution (Merck 1.02419.0250) for 10 mins. The sections were incubated in 60% isopropanol for 30 sec, rinsed with distilled water, counterstained with hematoxylin (Thermofisher Scientific, 7211) histological staining for 2 min, rinsed with distilled water, and mounted with Aquatex mounting media (Merck 1.08562.0050). Stained sections were imaged using Zeiss Imager.Z2 upright ApoTome, with Axiocam 512 color camera, at 10x, 40x, and 63x magnifications. For each section, at least three images were acquired. Imaging conditions were kept constant when acquiring images to be compared.

### Cell culture and glucose-sensing

HEK293T cells (ATCC, RRID: CVCL_0063) cells were cultured in Dulbecco′s Modified Eagle′s Medium (DMEM, Thermo Fisher Scientific, REF: 41965-039) containing 10% FBS (gibco, REF: A3160801) and 1% Penicillin/Streptomycin (P/S, Thermo Fisher Scientific, REF: 15140122). Glucose Sensing was performed by incubation in 2.5mM glucose DMEM (Thermo Fisher Scientific, REF: 11966-0259) + FBS and P/S for 6 hours and following incubation in 2.5mM or 25mM glucose DMEM + FBS and P/S for 24 hours. Prior harvest cells were washed with ice cold PBS.

### Transfection

One day before transfection cells were seeded at a concentration of 12000 cells/well in Poly-L-Lysine (45µl/well, 2.5µg/ml for 30min @ room temperature; Bio&SELL, REF: BS.L 7250) coated 96 well plates (Greiner Bio-One, Ref: 655-180). Co-transfection of pCMV4-Mlxipl-5’FLAG (20ng/well) and pCMV4-Mlx (25ng/well) isoforms was carried out overnight at a cell confluence of 60% using Lipofectamine 2000 (Thermo Fisher Scientific, REF: 11668019) according to the manufacturer’s protocol.

### Immunoblotting

Cells were treated with lysis buffer (1:1 = 20% SDS: Non-Reducing Lane Marker Sample Buffer (Thermo Fisher Scientific, REF: 39001) + beta-Mercaptoethanol) and denatured at 95°C for 10 min. Proteins were separated by SDS-PAGE and transferred to PVDF membranes by Western Blot overnight. Membranes were blocked in 4% milk. Anti-FLAG M2-Peroxidase (Sigma-Aldrich, REF: A8592, Lot: SLBK9652V), anti-beta-Actin (Santa Cruz, REF: sc-47778, Lot: #H1914), & anti-ChREBP (Novus Biologicals, REF: NB400-135, Lot: S2) were used as primary antibody and subsequently a corresponding secondary antibody when necessary. After incubation in chemiluminescent substrate (Thermo Fisher Scientific), bands were detected on a ChemiDocTM Imaging System (Bio-Rad).

### qPCR

RNA was extracted from cells using Total RNA Kit (peqGOLD, REF: 12-6834-02). After determination of RNA concentration per NanoDrop® Spectrophotometer ND-1000, cDNA was synthesized from equal RNA amounts using M-MLV Reverse Transcriptase (Promega, REF: M1708) followed by quantitative PCR with Takyon™ No ROX SYBR 2X MasterMix blue dTTP. Data analysis was performed using standard curve calculation and mRNA expression was normalized to human HPRT followed by normalization to one of the conditions: swift long isoform at 2.5mM glucose. Primer sequences are listed in (table S15).

### Luciferase Assay

Transcriptional activity was assessed using a Luciferase reporter driven by acetyl-CoA carboxylase (ACC1) Carbohydrate Response Element (ChoRE) (Tsatsos and Towle. Biochem Biophys Res Commun. 2006). The Dual-Luciferase® Reporter Assay System (Promega, REF: E1960) was used to assess transcriptional activity. The respective pCMV4-MLXIPL-5’FLAG isoform (20ng/well) was cotransfected with the corresponding pCMV4-MLX isoform (25ng/well), pGL3-ACC1-ChoRE firefly luciferase (100 ng/well) as well as pRL-Renilla luciferase (0.4ng/well) as internal control. Luminometry was performed as outlined in the manufacturer’s manual using Mithras LB 940 (Berthold technologies). Results were normalized to Renilla luciferase activity followed by normalization to one of the conditions: swift long isoform at 2.5mM glucose.

## Acknowledgments

We thank the Macaulay Library at the Cornell Lab of Ornithology for providing bird images. We also thank Elena Gracheva, Madeleine Junkins, Paul Sunnucks, and Jane Reznick for comments on the manuscript, and Emily Wheeler for editorial assistance.

## Authors Contribution

E.O., M-C.K., M.W.B., M.H., and T.B.S. conceptualized the study, did the analysis and the experiments, and wrote the manuscript. K.M.P. and M.S. lead and performed the MLXIPL experiments. S.Y.W.S., T.B., S.W., M.P., M-B.M., F.T-L., C.W., O.C., R.O., P.E., H.K., A.B., performed sampled preparation, sequencing and assembly of genomes. J.J. extracted tissue-specific RNA; S.W. did histological sectioning and staining; D.M.A assisted with annotation; A.G., K.S., G.W.L., A.M., N.B., N.M.A., L.M., A.v.B., A.R-G., D.L.A., W.A.B., contributed to sample collection; S.V.E., C.F-V., M.G. contributed resources or support facilitating genome sequencing.

## Data, Code, and Materials Availability

Genome assemblies are available on NCBI under the following accessions: *Acanthiza pusilla, Phylidonyris novaehollandiae, Furnarius rufus*, and *Trichoglossus moluccanus* are: PRJNA623276; *Florisuga fusca*: PRJNA622897; *Nymphicus hollandicus*: PRJNA819251; *Loriculus galgulus*: PRJNA1112892; *Ptilotula penicillata*: PRJNA1112893. Transcriptomic data are available on NCBI under the following accessions: *Calypte anna*: PRJNA1031337; *Apus apus*: PRJNA1031644; *Taeniopygia guttata*: PRJNA1032078; *Nymphicus hollandicus*: PRJNA1032113; *Phylidonyris novaehollandiae*: PRJNA1051659; *Trichoglossus moluccanus*: PRJNA1051684. The genome annotations, ortholog alignments, phylogenetic tree, aBSREL, PhyloAcc, kallisto, DESeq2 analysis result tables, as well as other files generated in this study from the raw data are available at DOI:<u>10.5281/zenodo.17403725</u>. TOGA annotations are available at https://genome.senckenberg.de/download/TOGA/. The scripts used in this study are available at https://github.com/maggieMCKO/nectargenomics and permanently archived (*171*). No new materials were generated for this study.

