## Supplementary Figures for "Convergent and lineage-specific genomic changes shape adaptations in sugar-consuming birds"

Science
AAAS

### Supplementary Materials for

#### Shared and lineage-specific genomic changes contribute to adaptations in sugar-consuming birds

**Authors:** Ekaterina Osipova[1,2,3,4*], Meng-Ching Ko[5*], Konstantin M. Petricek[6], Simon Yung Wa Sin[7,8,9], Thomas Brown[10,11], Sylke Winkler[10], Martin Pippel[12,13], Julia Jarrells[10], Susanne Weiche[14], Mai-Britt Mosbech[15], Fanny Taborsak-Lines[16,17], Chuan Wang[18,19], Orlando Contreras-Lopez[16,19], Remi-Andre Olsen[20,19], Philip Ewels[20,21], Daniel Mendez-Aranda[5,22], Andrea Gaede[23], Keren Sadanandan[5], Gabriel Weijie Low[5,24], Amanda Monte[25], Ninon Ballerstaedt[26], Nicolas M. Adreani[27,28], Lucia Mentesana[27], Auguste von Bayern[29], Alejandro Rico-Guevara[30,31], Scott V. Edwards[8,9], Carolina Frankl-Vilches[25], Heiner Kuhl[25,34], Antje Bakker[25], Manfred Gahr[25], Douglas L. Altshuler[32], William A. Buttemer[33] , Michael Schupp[6], Maude W. Baldwin[5†], Michael Hiller[2,3,4†], Timothy B. Sackton[1†]

* co-first authors

**The PDF file includes:**

Figs. S1 to S12
Tables S1 to S15

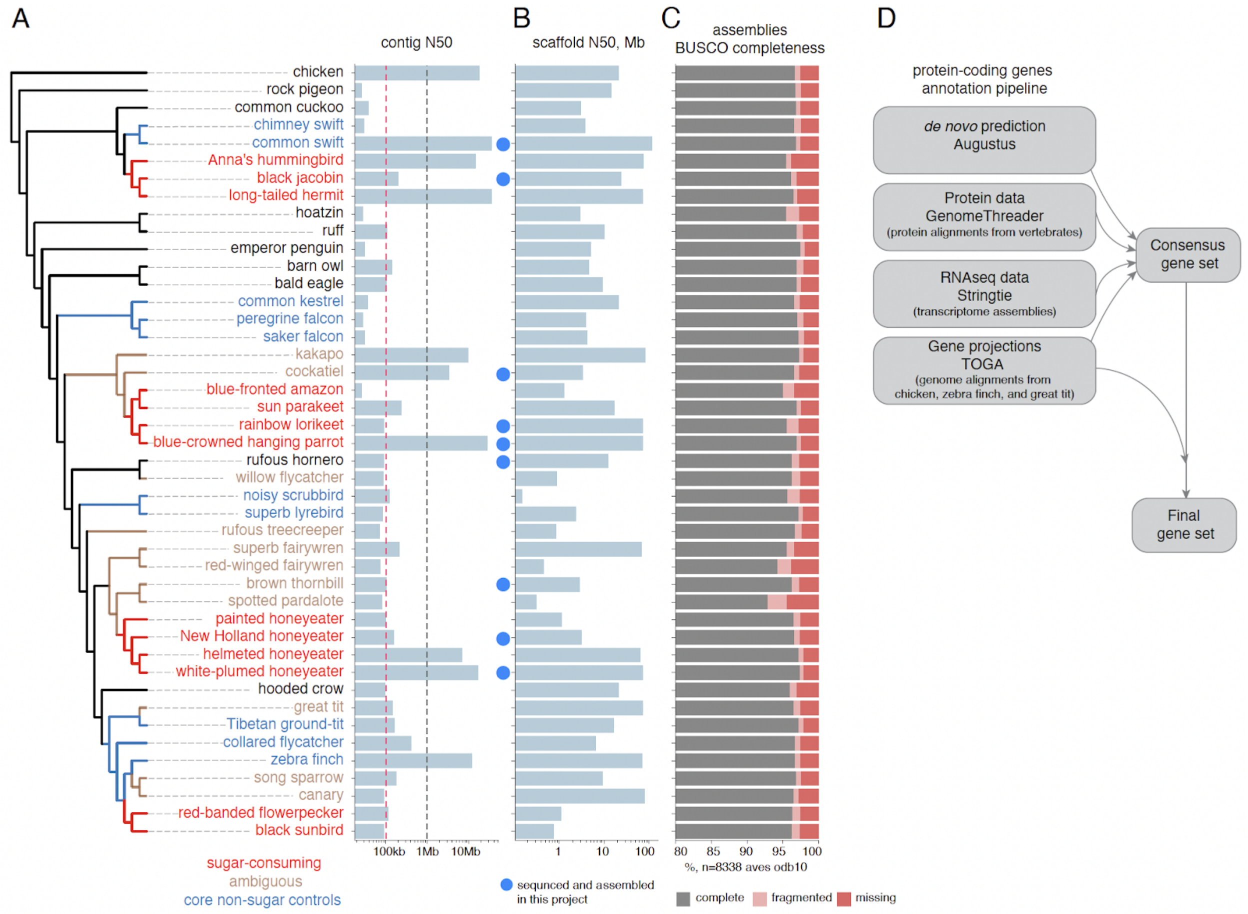


**Fig. S1. Assemblies and annotations**

(A) Phylogeny and contig N50 of the assemblies included in the study. Based on the dietary sugar intake, we classified species as core sugar-consuming (red font), core non-sugar controls (blue), or ambiguous (brown). Core non-sugar control sets were defined as groups phylogenetically close to sugar-consuming groups and comprising a similar number of species, but not taking any sugar in the diet. These were used as direct controls in the downstream analyses. Lineages that consume some fruit or nectar (but less than 50% of diet) or for which the ancestral state of sugar consumption is uncertain are defined as ambiguous (see Materials and Methods for details).

(B) Scaffold N50 of the assemblies included in the study.

(C) Genome BUSCO completeness of 8338 universal single-copy avian orthologs (BUSCO odb10 gene set).

(D) Protein-coding gene annotation pipeline used in the project.

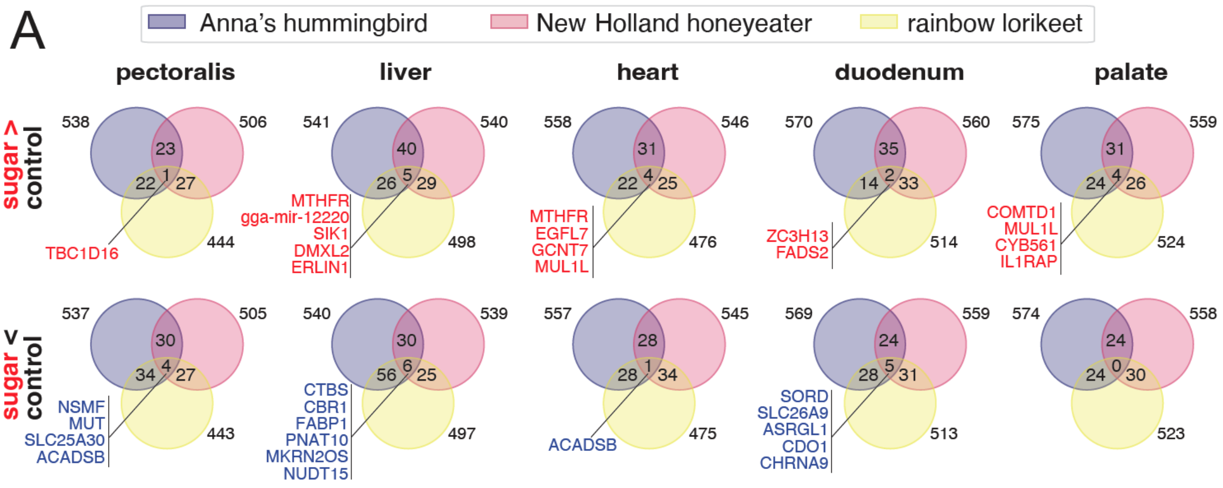
A
Anna's hummingbird
New Holland honeyeater
rainbow lorikeet
pectoralis
liver
heart
duodenum
palate
sugar > control
sugar < control
538
506
23
22
1
27
444
TBC1D16
541
540
40
26
5
29
498
MTHFR
gga-mir-12220
SIK1
DMXL2
ERLIN1
558
546
31
22
4
25
476
MTHFR
EGFL7
GCNT7
MUL1L
570
560
35
14
2
33
514
ZC3H13
FADS2
575
559
31
24
4
26
524
COMTD1
MUL1L
CYB561
IL1RAP
537
505
30
34
4
27
443
NSMF
MUT
SLC25A30
ACADSB
540
539
30
56
6
25
497
CTBS
CBR1
FABP1
PNAT10
MKRN2OS
NUDT15
557
545
28
28
1
34
475
ACADSB
569
559
24
28
5
31
513
SORD
SLC26A9
ASRGL1
CDO1
CHRNA9
574
558
24
24
0
30
523

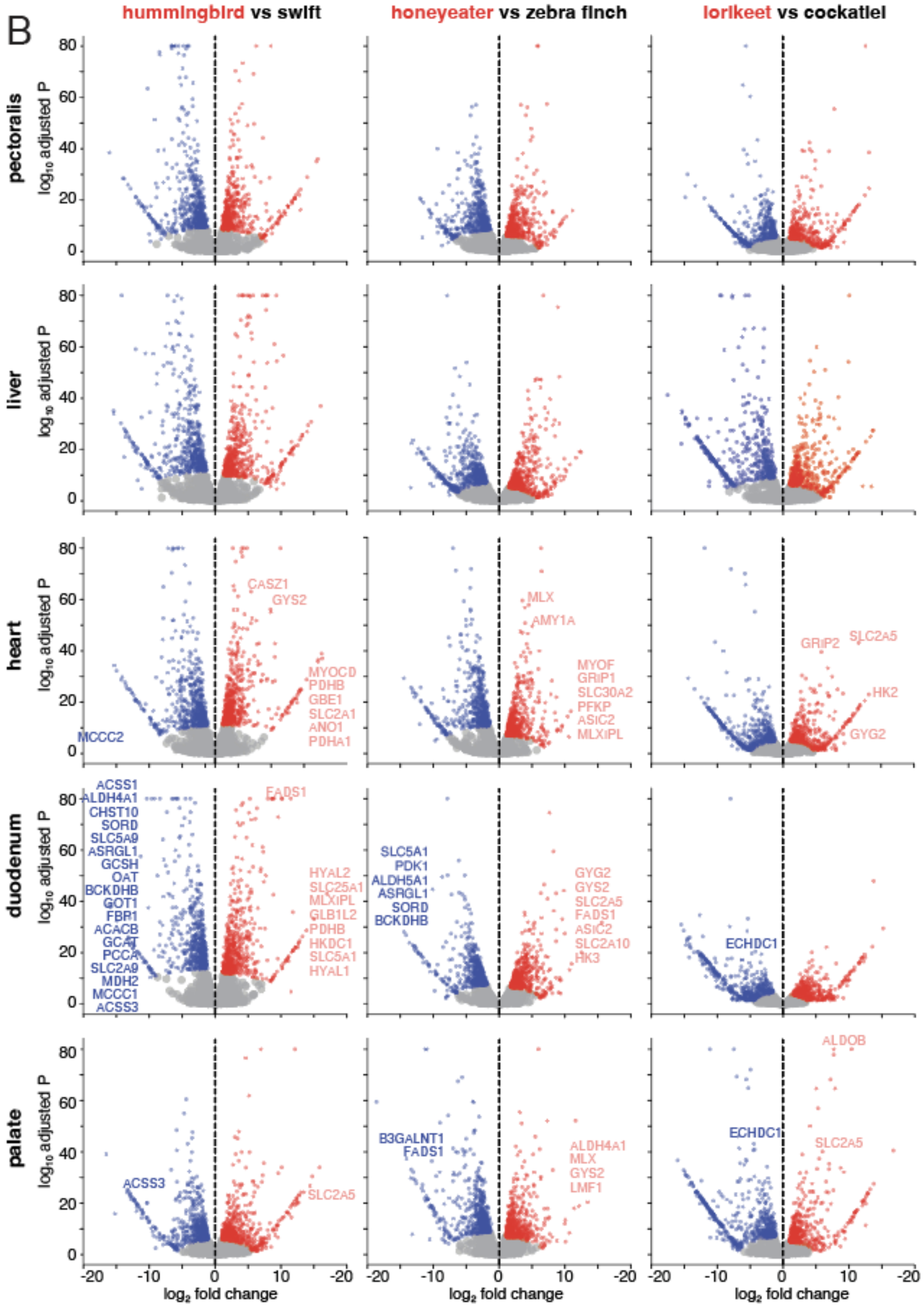
B
hummingbird vs swift
honeyeater vs zebra finch
lorikeet vs cockatiel
pectoralis
liver
heart
duodenum
palate
log10 adjusted P
log2 fold change
CASZ1
GYS2
MYOCD
PDHB
GBE1
SLC2A1
ANO1
PDHA1
MCCC2
MLX
AMY1A
MYOF
GRIP1
SLC30A2
PFKP
ASIC2
MLXIPL
GRIP2
SLC2A5
HK2
GYG2
ACSS1
ALDH4A1
CHST10
SORD
SLC5A9
ASRGL1
GCSH
OAT
BCKDHB
GOT1
FBP1
ACACB
GCAT
PCCA
SLC2A9
MDH2
MCCC1
ACSS3
FADS1
HYAL2
SLC25A1
MLXIPL
GLB1L2
PDHB
HKDC1
SLC5A1
HYAL1
SLC5A1
PDK1
ALDH5A1
ASRGL1
SORD
BCKDHB
GYG2
GYS2
SLC2A5
FADS1
ASIC2
SLC2A10
HK3
ECHDC1
ACSS3
SLC2A5
B3GALNT1
FADS1
ALDH4A1
MLX
GYS2
LMF1
ECHDC1
ALDOB
SLC2A5

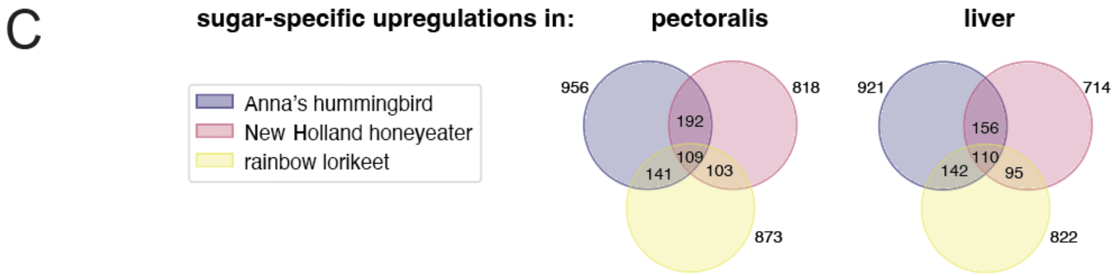


**Fig. S2. RNA-seq analysis**
(A) Venn diagrams presenting overlap between the gene sets of top 5% up- (top) or downregulated (bottom) genes in each of the five tested tissues in three sugar-feeders. Full lists of differentially expressed genes are in table S10.
(B) Volcano plots for five tested tissues showing genes with large expression changes in sugar-feeders. Red color represents upregulations (top 5%) in tissues of a sugar-consuming species, blue - downregulations (top 5%).
(C) Venn diagrams presenting overlap between sets of genes upregulated in pectoralis of sugar-feeders compared to liver (left) and in liver compared to pectoralis (right). Full lists of differentially expressed genes in the cross-tissue analysis are in table S13.

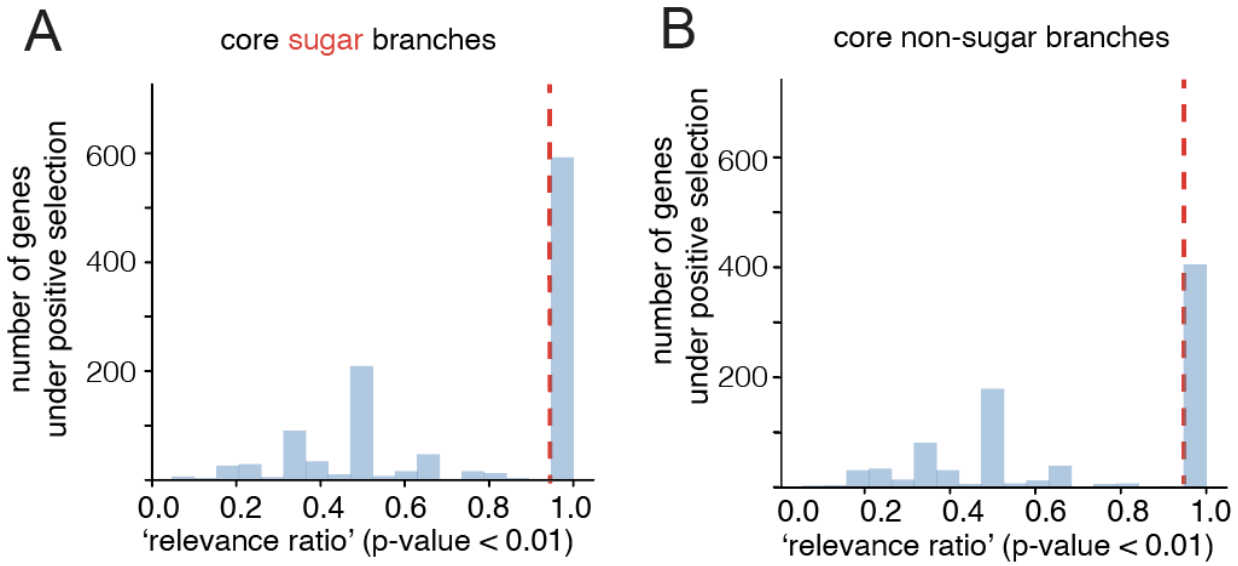


**Fig. S3. aBSREL analysis**
(A, B) Distribution of relevance ratio (the number of target branches showing a signal of positive selection divided by the total number of branches showing a signal of positive selection) for core sugar branches (A) and core non-sugar branches (B).

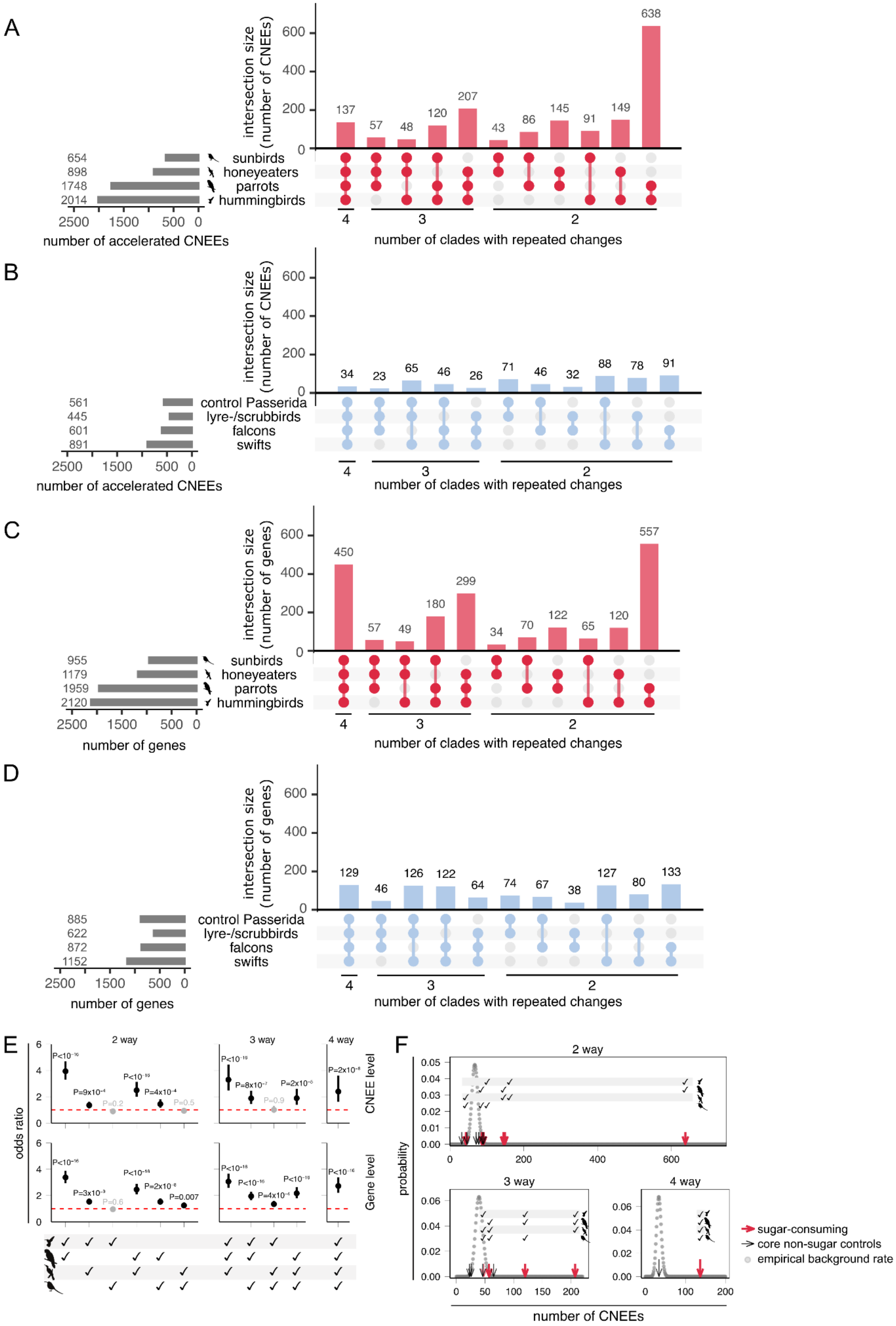
A
intersection size (number of CNEEs)
638
137
57
48
120
207
43
86
145
91
149
654
898
1748
2014
sunbirds
honeyeaters
parrots
hummingbirds
2500 1500 500 0
number of accelerated CNEEs
4
3
2
number of clades with repeated changes
B
intersection size (number of CNEEs)
34
23
65
46
26
71
46
32
88
78
91
561
445
601
891
control Passerida
lyre-/scrubbirds
falcons
swifts
2500 1500 500 0
number of accelerated CNEEs
4
3
2
number of clades with repeated changes
C
intersection size (number of genes)
450
57
49
180
299
34
70
122
65
120
557
955
1179
1959
2120
sunbirds
honeyeaters
parrots
hummingbirds
2500 1500 500 0
number of genes
4
3
2
number of clades with repeated changes
D
intersection size (number of genes)
129
46
126
122
64
74
67
38
127
80
133
885
622
872
1152
control Passerida
lyre-/scrubbirds
falcons
swifts
2500 1500 500 0
number of genes
4
3
2
number of clades with repeated changes
E
odds ratio
2 way
3 way
4 way
CNEE level
Gene level
P=0.007
F
2 way
3 way
4 way
probability
number of CNEEs
sugar-consuming
core non-sugar controls
empirical background rate

**Fig. S4. Comparison of accelerated CNEEs and associated genes in sugar-feeders versus their paired core non-sugar controls.**
(A-B) The upset plots in panels (A) for sugar-feeders and (B) for core non-sugar controls (corresponding outgroups for target sugar clades) display the count of repeatedly accelerated CNEEs in four-way, three-way, or two-way combinations. The horizontal bar plots on the left show the total number of accelerated CNEEs for each group.
(C-D) The upset plots in panels (C) for sugar-feeders and (D) for core non-sugar controls illustrate the count of genes associated with accelerated CNEEs in four-way, three-way, or two-way combinations. The horizontal bar plots on the left show the number of genes associated with accelerated CNEEs for each clade.
(E) Excess of repeated acceleration among sugar-feeders at CNEE (top) and gene levels (middle), showing more frequent acceleration in two-, three-, and four-way comparisons than in paired non-sugar controls. Significant differences (Fisher's exact test, two-sided) are shown in black.
(F) A binomial test evaluates the excess of molecular repeatability in target lineages (compared to background), by comparing the count of repeatedly accelerated CNEEs in sugar-consuming birds against the empirical background rate of accelerated CNEEs estimated from the average count in core non-sugar control birds. Downward-pointing arrows indicate the counts for each clade combination, red for sugar-feeders and black for core non-sugar controls. Bold arrows indicate significantly higher repeatability than empirical background distributions. All combinations involving sugar-consuming clades are significantly greater than those involving core non-sugar controls.

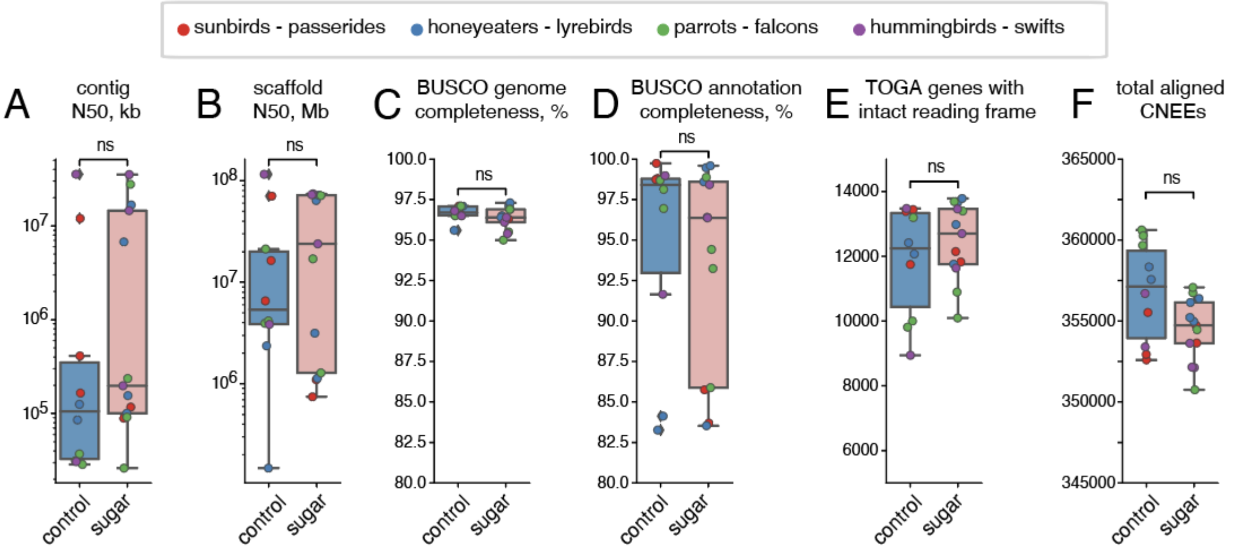


**Fig. S5. Comparison of genome assembly quality between sugar-consuming birds and controls.**
(A, B) contig (A) and scaffold N50 (B) value.
(C, D) Percent of BUSCO genes that are completely detected in the genome assembly (C) and gene annotation (D, BUSCO in annotation mode).
(E) Number of reference genes that have an intact reading frame in the TOGA annotation, which measures not only assembly completeness but also base accuracy.
(F) Total number of CNEEs aligned for each assembly.
Individual genomes are represented by dots, colored based on the taxonomic group. Box plots for each group indicate the first quartile, the median and the third quartile with whiskers extending up to 1.5 times the interquartile distance, showing outliers as data points that are 1.5 times the interquartile distance above the first or below the third quartile.
None of these metrics shows a significant difference between the two groups. The lowest p-value is 0.077 for CNEEs. Importantly, the three gene completeness metrics (C-E) are highly similar among the groups. Given the lack of a systematic difference in assembly quality or gene completeness, it is unlikely that differences in genome quality have a major impact on our comparative analysis.

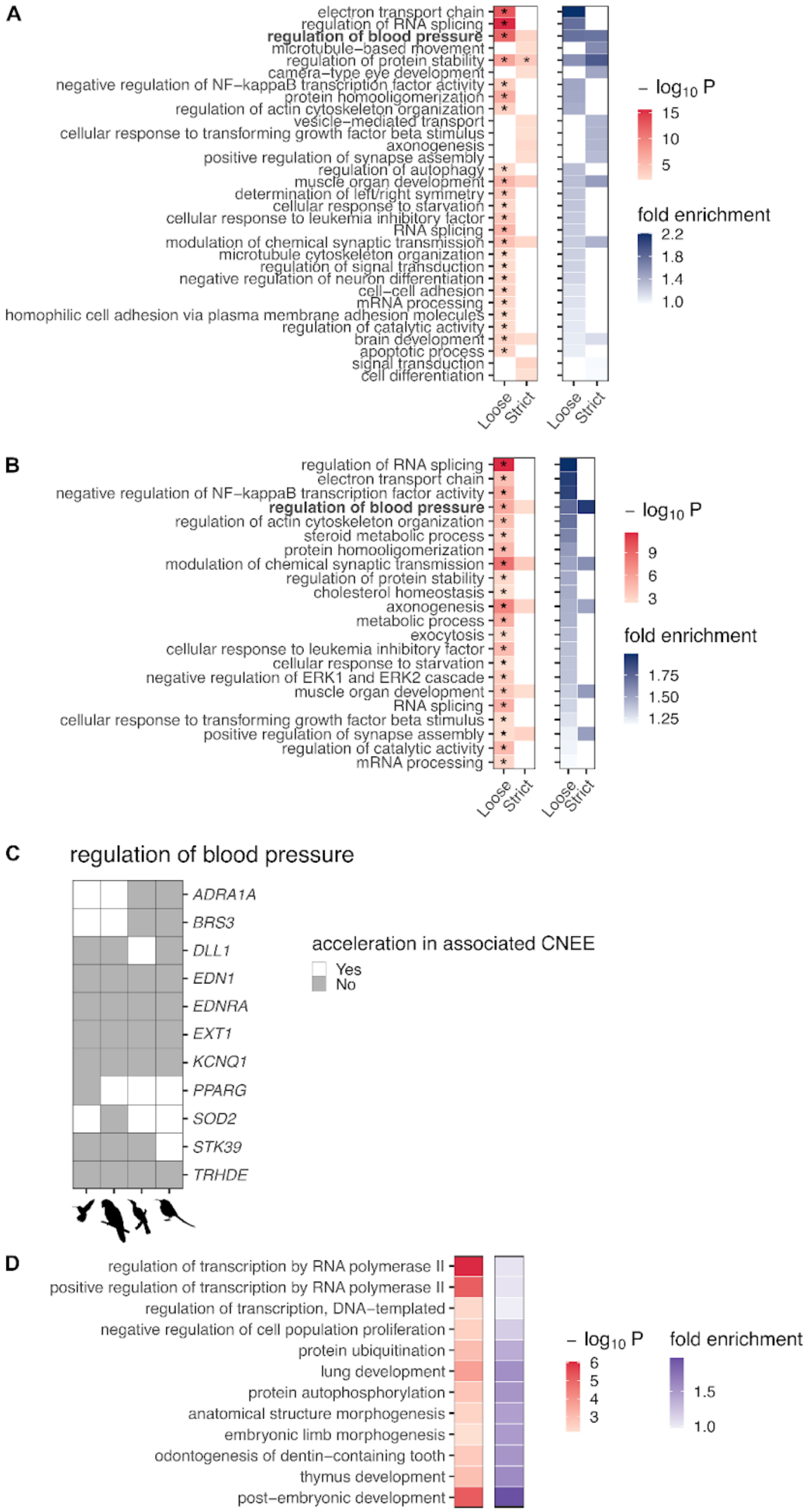
A
electron transport chain
regulation of RNA splicing
regulation of blood pressure
microtubule−based movement
regulation of protein stability
camera−type eye development
negative regulation of NF−kappaB transcription factor activity
protein homooligomerization
regulation of actin cytoskeleton organization
vesicle−mediated transport
cellular response to transforming growth factor beta stimulus
axonogenesis
positive regulation of synapse assembly
regulation of autophagy
muscle organ development
determination of left/right symmetry
cellular response to starvation
cellular response to leukemia inhibitory factor
RNA splicing
modulation of chemical synaptic transmission
microtubule cytoskeleton organization
regulation of signal transduction
negative regulation of neuron differentiation
cell−cell adhesion
mRNA processing
homophilic cell adhesion via plasma membrane adhesion molecules
regulation of catalytic activity
brain development
apoptotic process
signal transduction
cell differentiation
Loose
Strict
Loose
Strict
− log10 P
15
10
5
fold enrichment
2.2
1.8
1.4
1.0
B
regulation of RNA splicing
electron transport chain
negative regulation of NF−kappaB transcription factor activity
regulation of blood pressure
regulation of actin cytoskeleton organization
steroid metabolic process
protein homooligomerization
modulation of chemical synaptic transmission
regulation of protein stability
cholesterol homeostasis
axonogenesis
metabolic process
exocytosis
cellular response to leukemia inhibitory factor
cellular response to starvation
negative regulation of ERK1 and ERK2 cascade
muscle organ development
RNA splicing
cellular response to transforming growth factor beta stimulus
positive regulation of synapse assembly
regulation of catalytic activity
mRNA processing
Loose
Strict
Loose
Strict
− log10 P
9
6
3
fold enrichment
1.75
1.50
1.25
C
regulation of blood pressure
ADRA1A
BRS3
DLL1
EDN1
EDNRA
EXT1
KCNQ1
PPARG
SOD2
STK39
TRHDE
acceleration in associated CNEE
Yes
No
D
regulation of transcription by RNA polymerase II
positive regulation of transcription by RNA polymerase II
regulation of transcription, DNA−templated
negative regulation of cell population proliferation
protein ubiquitination
lung development
protein autophosphorylation
anatomical structure morphogenesis
embryonic limb morphogenesis
odontogenesis of dentin−containing tooth
thymus development
post−embryonic development
− log10 P
6
5
4
3
fold enrichment
1.5
1.0

**Fig. S6. Limited shared enrichments in non-sugar controls and lack of metabolism-related pathways contrasts with extensive metabolic adaptations in sugar-feeders.**

(A-B) GO enrichment analysis for genes near CNEEs accelerated in sugar-consuming clades. (A) shows results for the union set of CNEEs accelerated in any sugar-consuming clade, while (B) shows results for CNEEs repeatedly accelerated in at least two clades. Heatmap shading represents p-values (red) from hypergeometric tests and corresponding fold enrichment values (purple). Asterisks (*) indicate FDR-adjusted p < 0.05. Only GO terms with FDR-adjusted p < 0.3 are shown. For both panels, although the strict criterion yielded fewer significantly enriched GO terms compared to the loose criterion, fold enrichment values substantially increased for terms that remained enriched, indicating stronger signal robustness when applying more stringent thresholds.

(C) Genes associated with sugar-specific accelerated CNEEs in regulation of blood pressure. Gray-filled cells indicate the presence of accelerated CNEEs near the corresponding gene in each sugar-consuming clade.

(D) GO enrichment analysis for genes near CNEEs repeatedly accelerated in at least two non-sugar control clades. Heatmap shading represents p-values (red) from hypergeometric tests and corresponding fold enrichment values (purple). Only GO terms with FDR-adjusted p < 0.3 are shown.

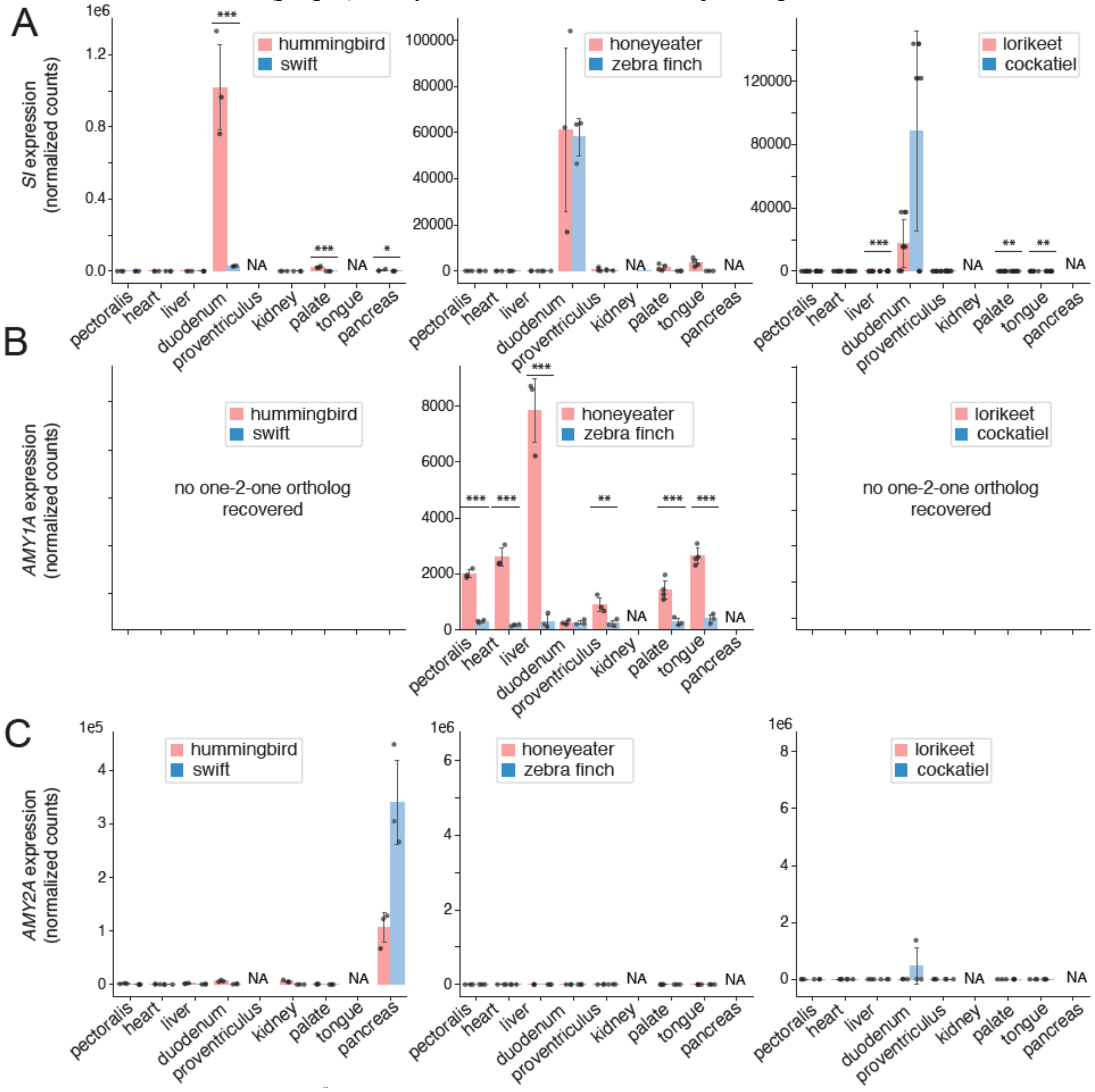

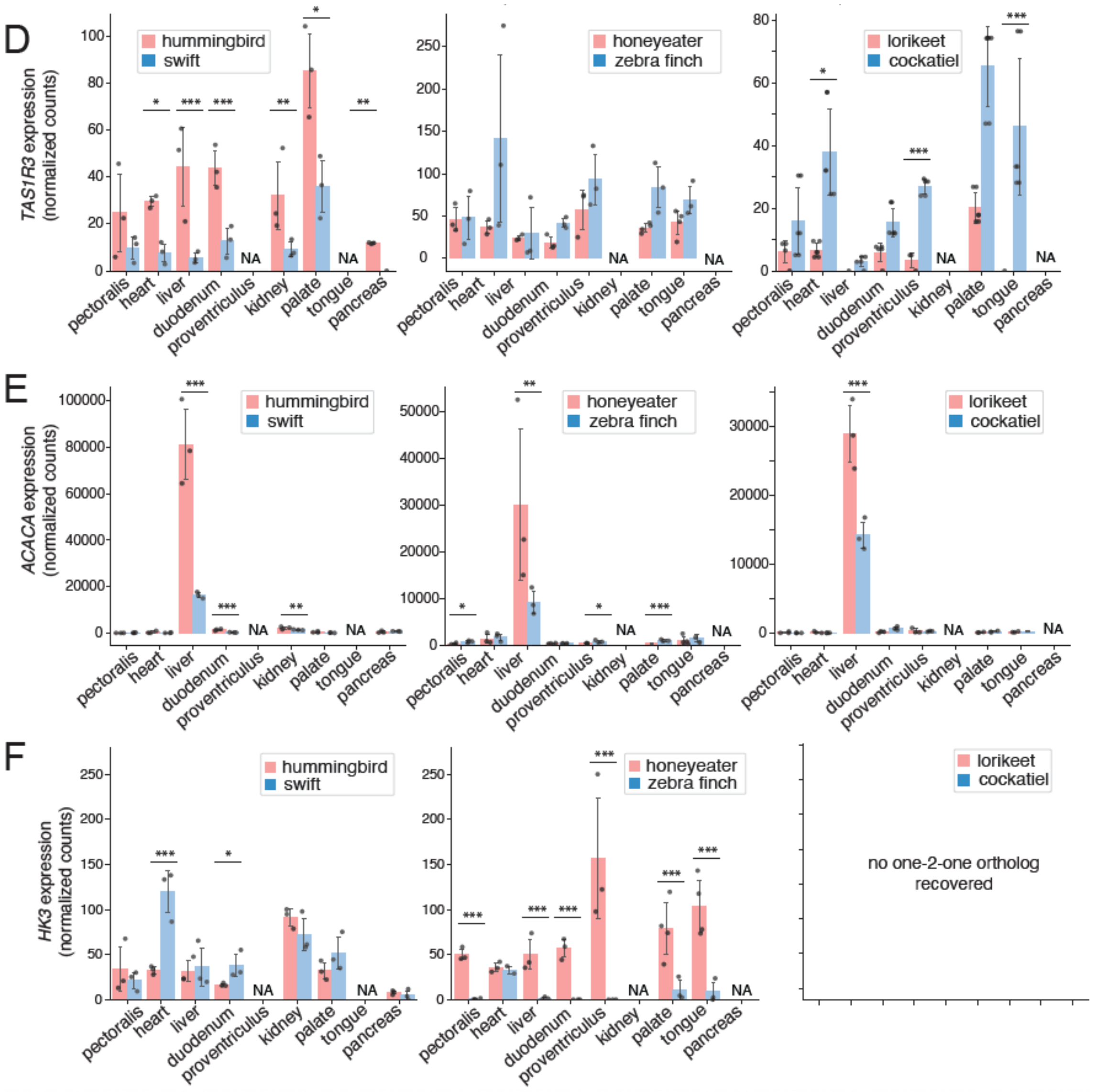


**Fig. S7. Candidate gene expression across all tissues in all pairs of sugar-control groups.**
(A) *SI* shows largely duodenum-specific expression across all pairs. *SI* ortholog was manually recovered in honeyeater.
(B) *AMY1A* is missing in the genomes of hummingbirds and many parrots, and appears to be uniformly upregulated across most sampled honeyeater tissues.
(C) *AMY2A* orthologs were recovered manually in all three pairs of species; *AMY2A* was not included in the initial DESeq2 analysis. *AMY2A* shows a repeated pattern of selection: it is associated with accelerated CNEEs in all four clades and shows pancreas-specific expression in hummingbirds (pancreas of the other two lineages were not sampled).
(D) *TAS1R3* in hummingbirds is most highly expressed in the palate (a likely location of taste buds) and is also upregulated in other tissues relative to the swift; upregulation was not observed in honeyeaters or lorikeets (cockatiels exhibited higher expression levels than lorikeets in many tissues).
(E) *ACACA* - the gene encoding the rate-limiting enzyme of the fatty acid biosynthesis pathway - shows a significant upregulation in the liver of all sugar-consuming birds.
(F) *HK3* shows honeyeater-specific upregulation across all tested tissues (except the heart). The *HK3* ortholog was not recovered in the lorikeet.
* signifies p-value < 0.05, ** - p-value < 0.01, and *** - p-value < 0.001 (FDR-corrected).

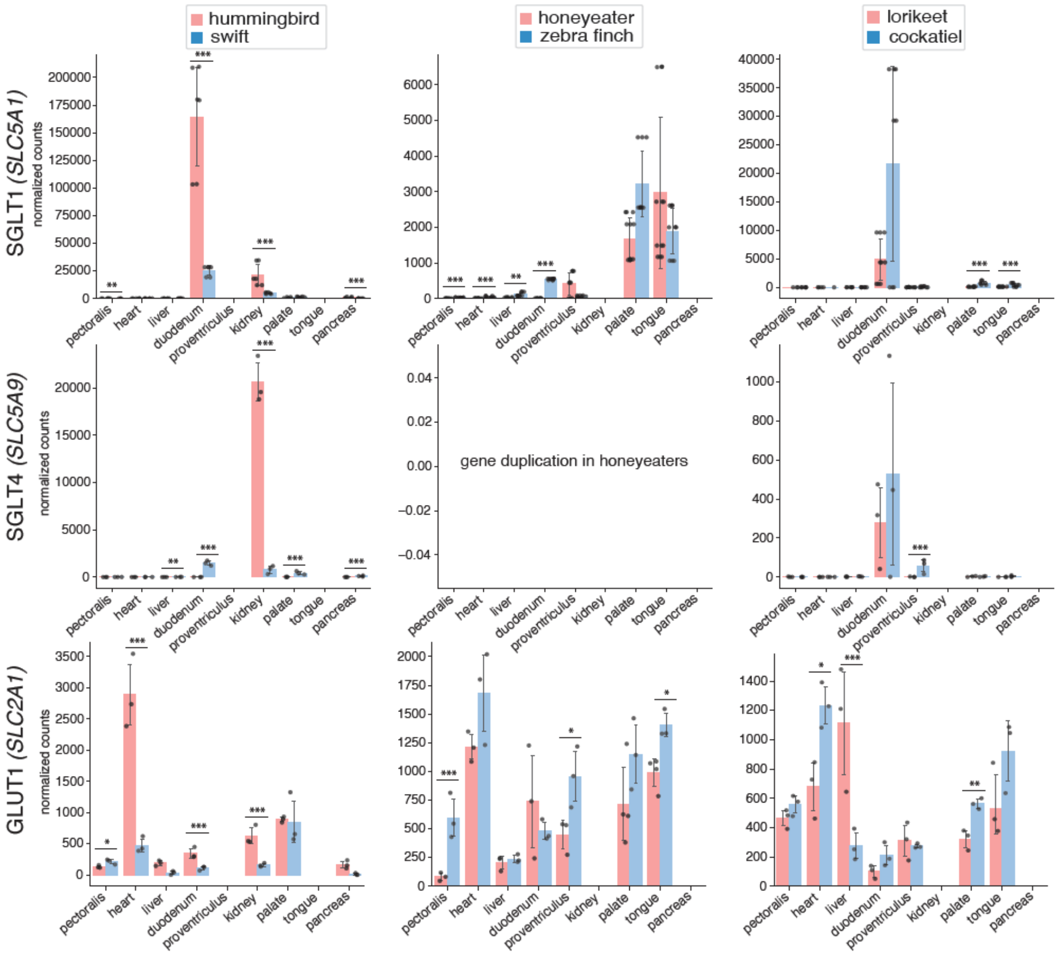

hummingbird
swift
honeyeater
zebra finch
lorikeet
cockatiel
SGLT1 (SLC5A1)
SGLT4 (SLC5A9)
GLUT1 (SLC2A1)
normalized counts
gene duplication in honeyeaters
pectoralis
heart
liver
duodenum
proventriculus
kidney
palate
tongue
pancreas

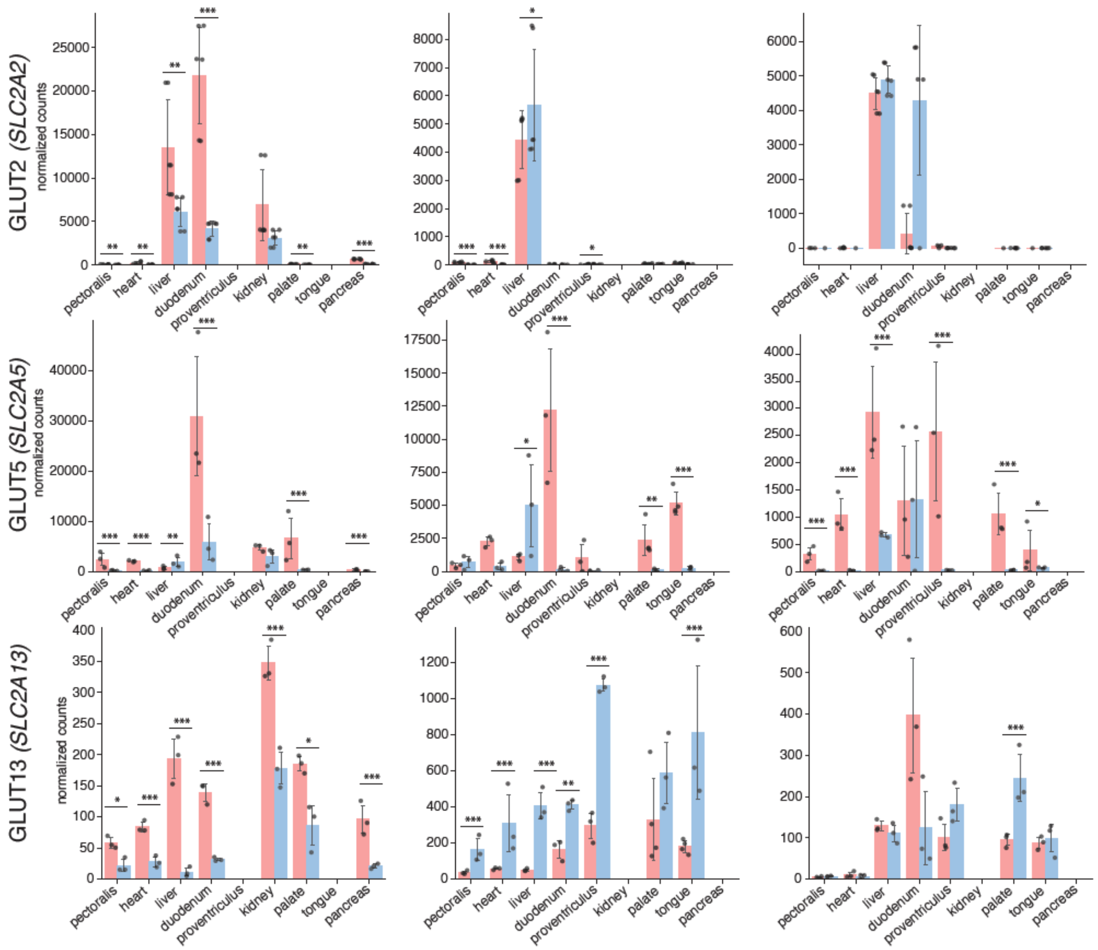


**Fig. S8. Gene expression of systemic sugar transporters across all tissues in all sugar-control comparison pairs.**

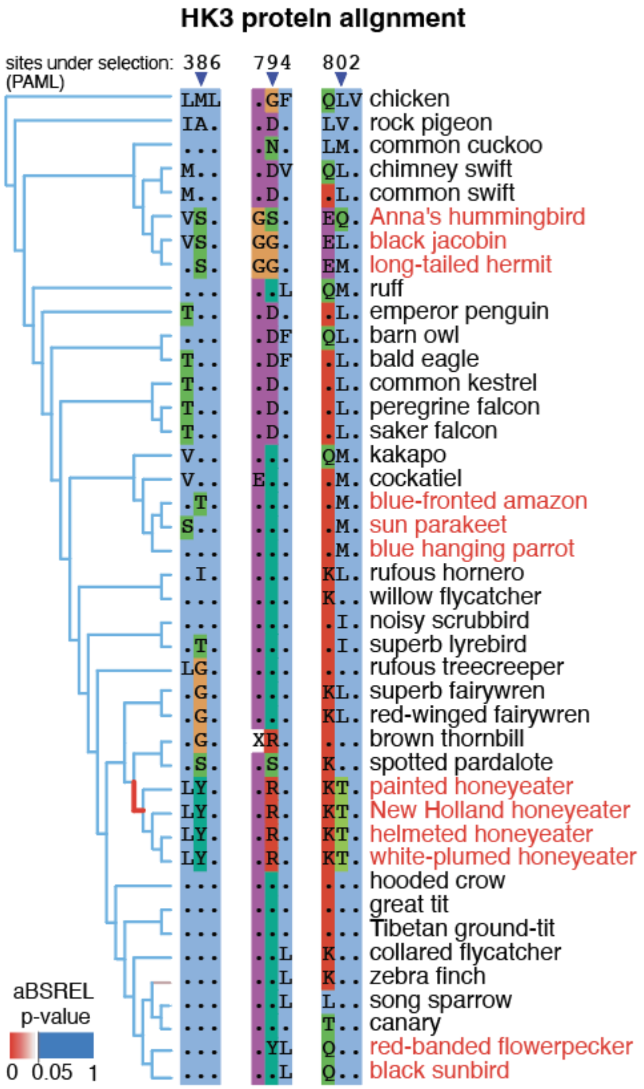


**Fig. S9. Positive selection in hexokinase 3 (HK3) in the ancestor of honeyeaters.**
Selected positions (PAML Bayes Empirical Bayes P>0.5) of the protein alignment of hexokinase 3 showing amino acid substitutions that occurred in the ancestral honeyeater branch. Branches are colored according to the aBSREL default model branch-corrected p-value.

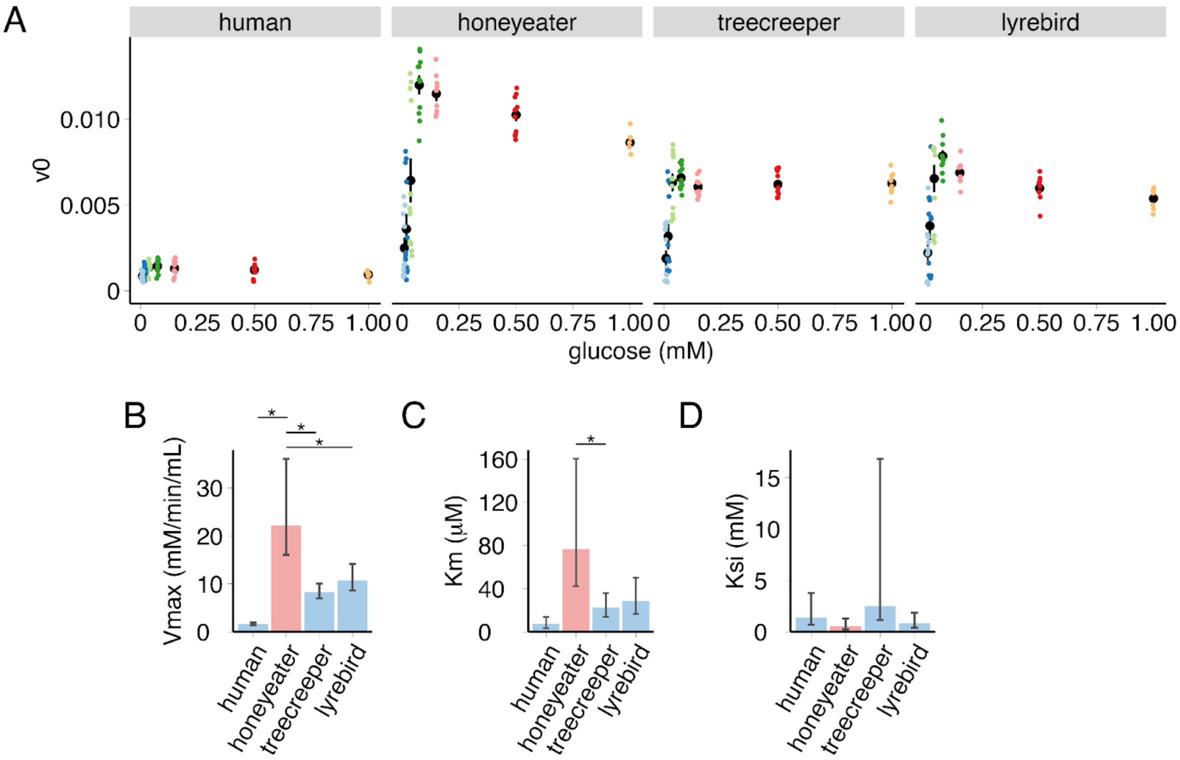


**Fig. S10. Kinetic properties of recombinant hexokinase 3 (HK3) from human, honeyeater, treecreeper, and lyrebird.**
(A) Initial velocity (v0) of HK3 enzymes at various glucose concentrations. Each point represents the estimated v0 of one sample. Sample sizes for each glucose concentration and enzyme combination range from 7 to 13.
(B) Maximum velocity (Vmax) of recombinant HK3 from the four species. Human: 1.6; honeyeater: 21.1; treecreeper: 8.0; lyrebird: 10.6 (mM/min/mL).
(C) Michaelis constant (Km) of recombinant HK3 from the four species. Human: 7.3; honeyeater: 65.9; treecreeper: 26.2; lyrebird: 30.6 (µM).
(D) Substrate inhibition constant (Ksi) of recombinant HK3 from the four species. Human: 1.5; honeyeater: 0.59; treecreeper: 3.1; lyrebird: 0.86 (mM). Error bars in panels B, C, and D indicate 95% confidence intervals.
In (B-D) * signifies p-value $< 0.05$.

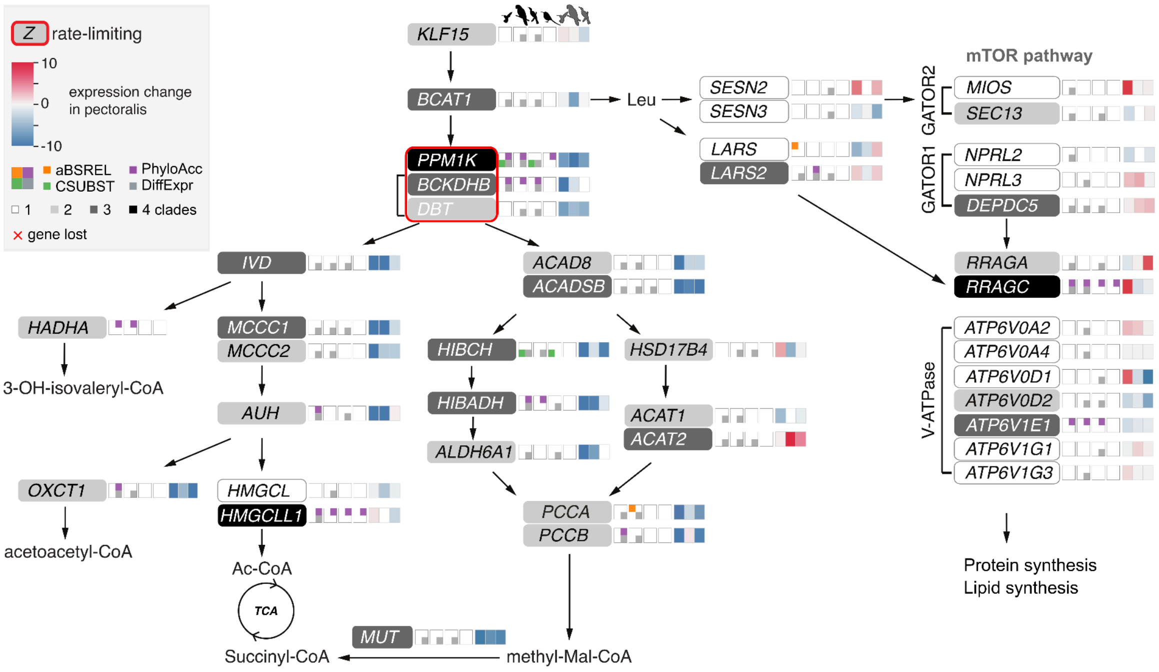


**Fig. S11. Genes of the branched-chain amino acid pathway are downregulated in sugar-feeders.** Large effects of regulatory evolution in branched-chain amino acid metabolism. Schematics as in Fig. 3B. The heatmap panels to the right of the quadrants illustrate the differential expression in the pectoralis of hummingbirds, parrots, and honeyeaters compared to their respective outgroups. Pathway modified from (87).

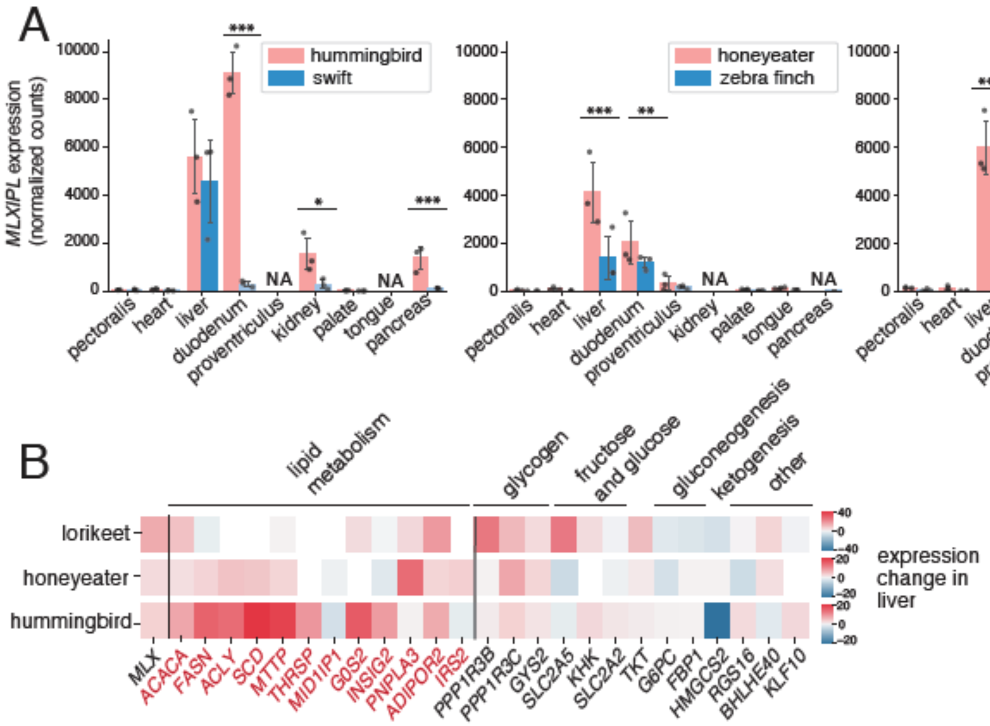

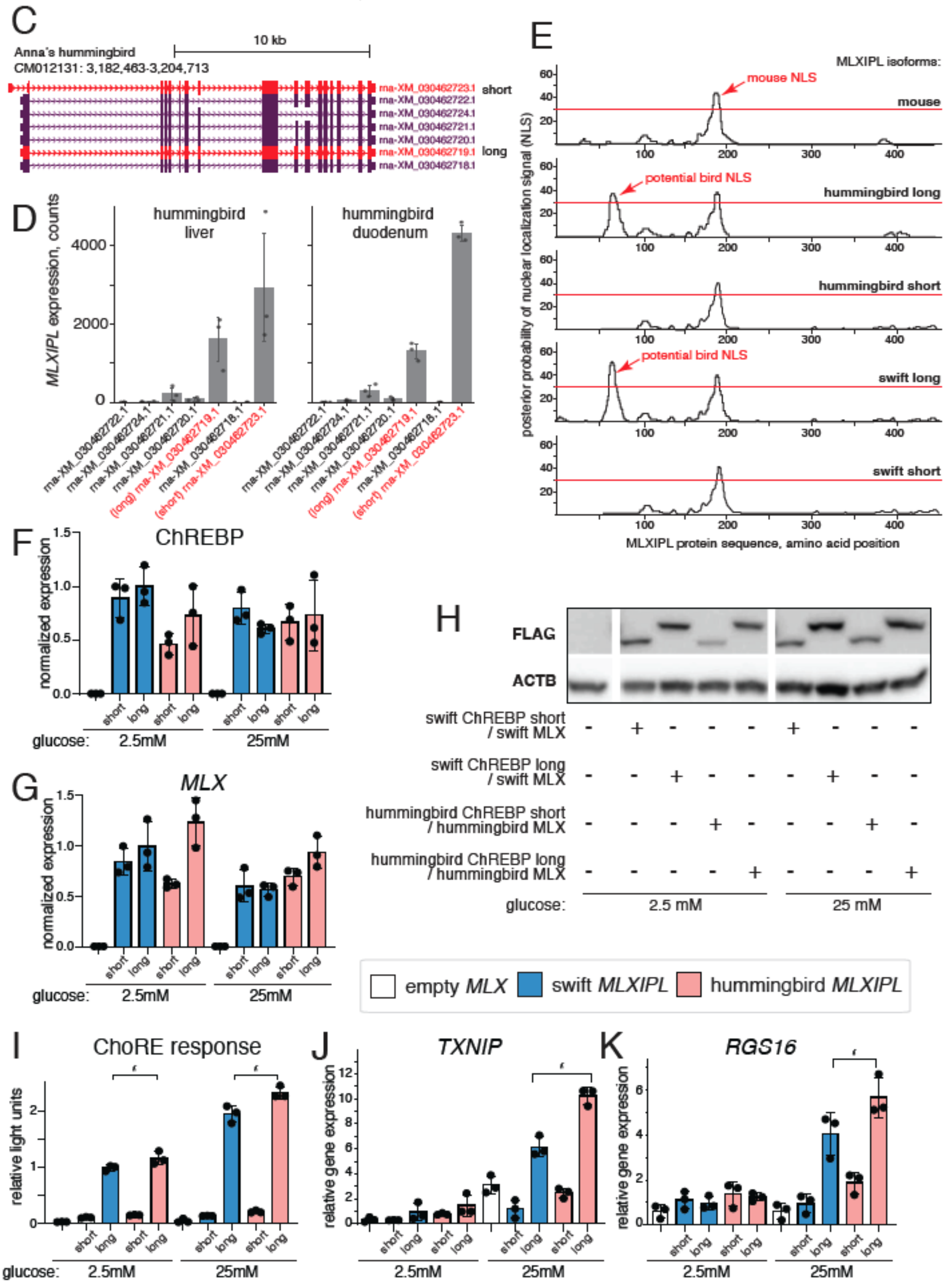


**Fig. S12. MLXIPL**

(A) *MLXIPL* expression in all tested tissues of sugar-consuming species compared to their corresponding outgroups.
(B) Expression changes of *MLXIPL* target genes in the liver.
(C) *MLXIPL* isoforms annotated by RefSeq for Anna's hummingbird, shown in the UCSC genome browser screen. Main isoforms (later referred to as short and long) are highlighted in red.
(D) Relative expression of all *MLXIPL* isoforms in liver and duodenum of the hummingbird. Main isoforms (later referred to as short and long) are highlighted in red.
(E) Theoretical prediction of nuclear-localization signals (NLS) in MLXIPL isoforms of mouse and two birds shows that the long bird isoform gains a new NLS, potentially making it the main functional isoform in birds.
HEK293T cells were transfected with depicted MLXIPL-FLAG and MLX isoforms and cultivated at 2.5mM or 25mM glucose for 24h.
(F, G) mRNA expression of MLXIPL and MLX isoforms.
(H) Immunoblot of ChREBP-FLAG protein with beta-Actin as loading control.
(I) Transcriptional activity of *MLXIPL* isoforms.
(J, K) Expression of two target genes of *MLXIPL* (*TXNIP* and *RGS16*).
In (A,F,G,I-K) * signifies p-value < 0.05, ** - p-value < 0.01, and *** - p-value < 0.001.
In (F,G, and I-K) relative light units and gene expression were normalized to one of the conditions: swift long isoform at 2.5mM glucose.

**Captions for Tables S1-S15**

Table S1: Information of the species and assemblies used in the analysis
Table S2: Genes under positive selection (relevance ratio = 1) per sugar-consuming group and control group
Table S3: GO enrichment analysis on the set of genes under selection in at least two sugar-consuming clades
Table S4: GO enrichment analysis on the set of genes under selection in at least one sugar-consuming clade
Table S5: GO enrichment analysis on the set of genes under selection in at least two non-sugar control clades
Table S6: GO enrichment of genes associated with CNEEs accelerated in any sugar-consuming clade
Table S7: GO enrichment of genes associated with CNEE-level repeated acceleration in at least two sugar-consuming clades
Table S8: Type 2 diabetes-associated genes with CNEE-level repeated acceleration in all four sugar-consuming clades
Table S9: Overview of samples collected for RNA-seq analysis
Table S10: Pairwise analysis: differentially expressed genes across all tested tissues
Table S11: Pathways enriched for upregulated genes pairwise differential gene expression analysis
Table S12: Pathways enriched for downregulated genes pairwise differential gene expression analysis
Table S13: Cross tissue analysis: genes differentially expressed between liver and pectoralis of sugar feeders and not in controls
Table S14: Pathways enriched for genes differentially expressed between liver and pectoralis of at least two sugar-feeders and not in controls
Table S15: Primers used for MLXIPL, MLX, and MLXIPL target genes
